# PRC2-binding regions are hotspots of DNA methylation entropy in pre-cancer and cancer

**DOI:** 10.64898/2026.10.11.758285

**Authors:** Djansel Bukovec, Monika Simjanoska Misheva, Marija Gjorgjievska, Goran Kungulovski

## Abstract

DNA methylation aberrations and variability are pervasive features of cancer, yet their relationship with chromatin regulation remains incompletely understood. Across hundreds of matched tumor–normal methylomes, regions with the highest methylation entropy in both tumor and normal tissues were consistently enriched for PRC2-associated factors. In tumors, inter-sample methylation variability was increased across the full range of PRC2-associated levels, while mean methylation progressively increased from low- to high-PRC2-binding regions; this gradient was substantially weaker in matched normal tissues. PRC2-associated regions with higher DNA methylation levels were preferentially associated with poised and repressed chromatin states, reduced chromatin accessibility, and reduced gene expression. Independent normal–precancer–cancer cohorts showed that this PRC2-associated methylation state emerges early, with precancerous and malignant tissues displaying highly similar profiles that are distinct from those of healthy tissues. Single-cell colon cancer methylomes independently confirmed increased methylation entropy at PRC2 regions. Higher PRC2-region entropy was associated with scores reflecting hypoxic states, aggressive tumor features, and poorer survival outcomes. Collectively, these findings identify PRC2-binding regions as hotspots of DNA methylation variability that become dysregulated during tumorigenesis.

## INTRODUCTION

Tumor development is accompanied by extensive genetic and epigenetic heterogeneity that contributes to variation in cellular identity, transcriptional state, and phenotypic behavior (1,2). Among epigenetic mechanisms, DNA methylation is a central regulator of gene expression and genome organization, and cancer-associated methylomes are classically characterized by focal CpG hypermethylation together with broad genomic hypomethylation (3,4). Large-scale profiling efforts, particularly through The Cancer Genome Atlas (TCGA), have established aberrant DNA methylation as a pervasive feature of human malignancies and have identified tumor subgroups with distinct methylation signatures across tissues (5–7). These observations have shifted the view of cancer beyond a purely genetic disease toward one in which disruption of epigenetic regulation is an integral component of malignant transformation and tumor evolution (8–11). In terms of DNA methylation, this disruption is not limited to differential changes in DNA methylation levels, but tumors also exhibit increased methylation variability within individual lesions and across patients and cancers (12–24).

These findings suggest that DNA methylation variability represents a distinct layer of cancer transformation emanating from impaired maintenance of epigenetic states and stochastic drift, potentially fostering phenotypic plasticity, microenvironmental adaptation, invasion, disease progression, and treatment resistance (1,14,25). A key unresolved question is whether DNA methylation variability accumulates randomly across the genome or preferentially localizes to specific regulatory environments. Findings from our previous work and independent studies demonstrated substantial crosstalk between DNA methylation and PRC2 in both cancer and ageing (5,26–35). PRC2 establishes repressive chromatin through H3K27me3 deposition and is central to developmental gene silencing, cellular identity, and epigenetic memory. These observations suggest that PRC2-regulated chromatin may represent an intrinsically vulnerable epigenetic compartment whose dysregulation is amplified during malignant transformation.

We systematically addressed this question using an unbiased analysis of transcription factor binding enrichment in high-entropy regions of hundreds of matched normal, pre-malignant, and malignant tissues from independent datasets, which consistently highlighted PRC2 members and associated factors across cancer types. We characterized this relationship across methylomes by integrating methylation entropy and inter-sample variability with PRC2 binding, chromatin state, accessibility, transcription, and clinical phenotypes.

## RESULTS

### High-entropy methylation regions are marked by PRC2-associated factors across the normal-to-cancer continuum

Expanding on our previous observation that DNA methylation entropy is increased in tumors relative to matched normal tissues (26), we next asked whether regions with the highest methylation entropy preferentially overlap specific transcription factor (TF) binding sites. As a discovery cohort, we analyzed paired Illumina HumanMethylation450 data from TCGA across 16 cancer types. We quantified DNA methylation entropy using the proportion of intermediately methylated sites (PIM) and Shannon entropy (SE), and tested highly entropic regions for enrichment of ENCODE 49 TF-binding peaks. Because tissue-matched TF ChIP-seq profiles were not uniformly available across all cancer types, we used binding-site annotations derived from human embryonic stem cells (hESC) as a common proxy across tissues. Across cancer types, PRC2 complex members and PRC2-associated factors were consistently represented among the ten most enriched factors at high-entropy regions in both tumor and corresponding normal tissues (**Figure 1**; **Supplementary Figures 1–2**).

**Figure 1.**
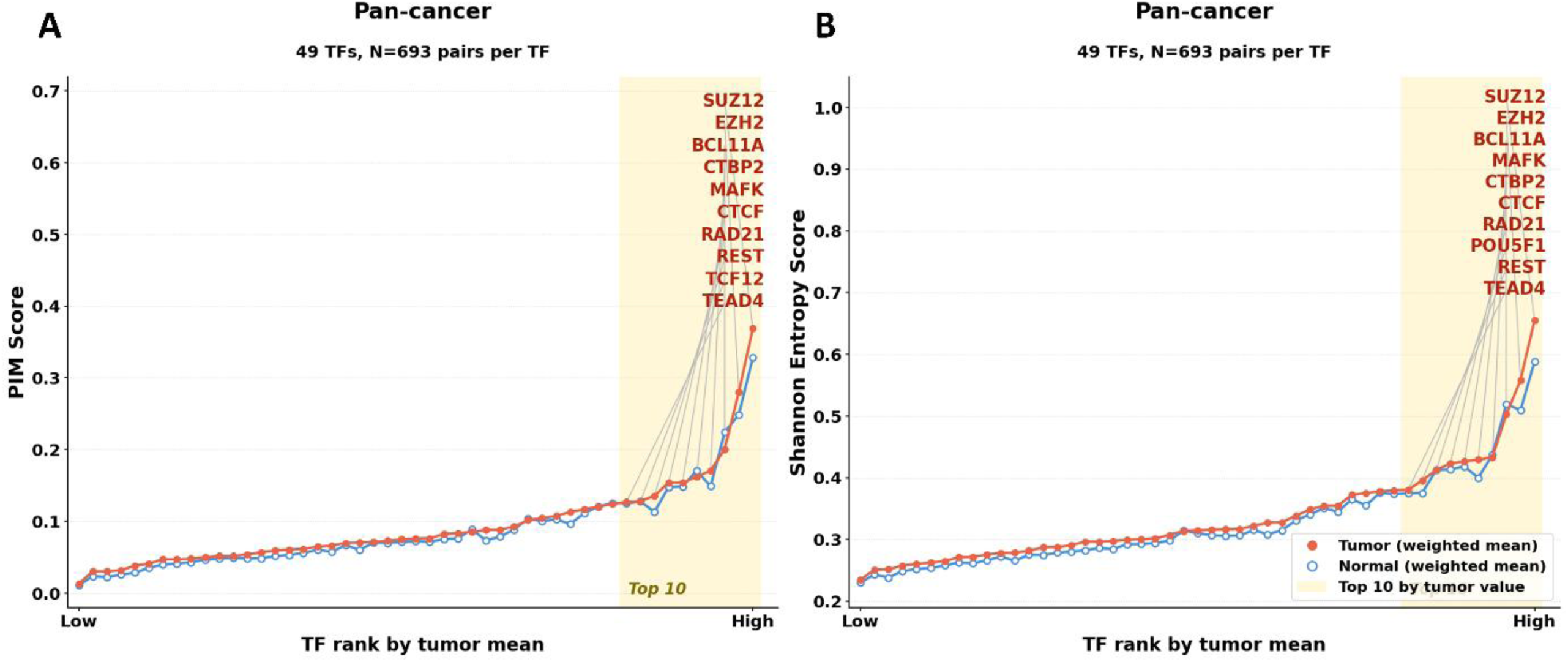
Pan-cancer DNA methylation disorder across transcription factor binding regions. Pan-cancer weighted mean DNA methylation variability was quantified within binding regions of 49 transcription factors (TFs) using 693 matched tumor–normal pairs per TF. TF-binding annotations were derived from human embryonic stem cell datasets and used as a common reference across cancer types. TFs are ordered from lowest to highest tumor-associated value. A) Proportion of intermediately methylated sites (PIM). B) Shannon entropy. Tumor values are shown as filled red circles and matched normal values as open blue circles. The shaded area denotes the 10 TFs with the highest tumor values in each analysis; corresponding TFs are labeled. PRC2 components, including SUZ12 and EZH2, ranked among the TF-associated regions with the highest PIM and Shannon entropy in both tumor and normal tissues

We subsequently examined 18 independent GEO DNA methylation cohorts totaling 3,110 samples of breast cancer or breast cancer and field effect (36–40), colorectal cancer progression (41–43), lung cancer and pre-invasive lung lesions (44,45), hepatocellular carcinoma (46), prostate cancer progression (47,48), esophageal cancer progression (49,50), gastric cancer progression (51), cholangiocarcinoma (52), and oral squamous cell carcinoma (53) spanning healthy donor tissues, tumor-matched adjacent normal tissues, adjacent tissues sampled at different distances from the tumor, precancerous lesions, invasive tumors, and progressive and regressive in situ lesions **(Supplementary Figures 3–8**). Despite differences in tissue type and disease stage, high-entropy regions in these datasets similarly showed preferential enrichment for PRC2 components or PRC2-associated factors.

To corroborate the association between DNA methylation entropy and PRC2-regulated regions, we quantified PIM and Shannon entropy within EZH2-binding sites and compared them with CTCF- and CHD1-binding regions, which were also identified as high-entropy candidates. For each factor, matched random genomic CpG regions with comparable genomic characteristics were included as controls. EZH2-binding regions showed significantly higher DNA methylation entropy than CTCF- and CHD1-binding regions and their corresponding matched random controls (**Figure 2**). The same pattern was reproduced in the independent validation datasets (**Supplementary Figure 9**). Consistent with our previous genome-wide analysis (26), both PIM and SE were consistently higher in tumors than in matched normal tissues (**Supplementary Figures 10 and 11**). The two metrics were strongly correlated (r = 0.86), indicating that PIM and SE capture largely overlapping patterns of methylation variability. Importantly, their distributions showed marked differences across cancer types, with some malignancies exhibiting consistently higher entropy than others, indicating that the magnitude of DNA methylation variability is strongly cancer-type specific (**Supplementary Figure 11C and F**). Within tumor samples, the levels of methylation entropy at EZH2-binding regions were significantly higher than at attribute-matched random regions in both the discovery and validation cohorts (**Supplementary Figures 12 and 13**).

**Figure 2.**
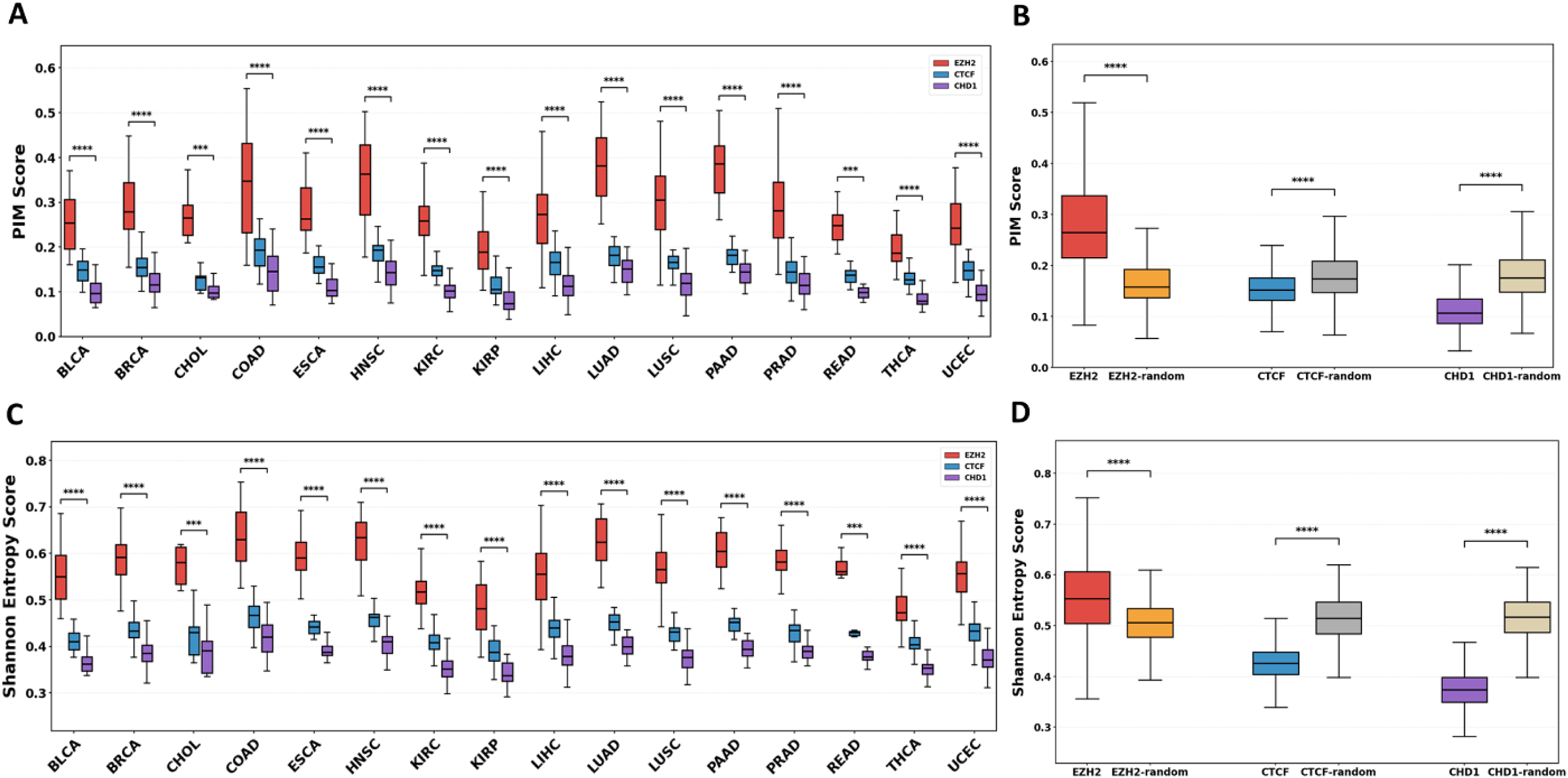
DNA methylation variability across EZH2-, CTCF-, and CHD1-binding regions and matched random controls. A) PIM scores across 16 cancer types within EZH2-, CTCF-, and CHD1-binding regions. B) Pan-cancer PIM scores within EZH2-, CTCF-, and CHD1-binding regions and their corresponding matched random control regions. C) Shannon entropy scores across 16 cancer types within EZH2-, CTCF-, and CHD1-binding regions. D) Pan-cancer Shannon entropy scores within EZH2-, CTCF-, and CHD1-binding regions and their corresponding matched random control regions. Boxplots show the median, interquartile range, and whiskers. Statistical significance is indicated above the corresponding comparisons (***P < 0.001; ****P < 0.0001).

To determine whether elevated methylation entropy at PRC2-associated regions reflects an intrinsic property of malignant cells rather than cellular admixture in bulk tissue, we analyzed single-cell DNA methylation data from colorectal cancer. PIM and Shannon entropy were benchmarked against the epiCHAOS metric in 509 cancer cells and 80 normal cells (54,55). Cancer cells showed clearly shifted distributions toward higher methylation variability across all three metrics compared with normal cells (**Figure 3**). PIM and SE were strongly correlated (r = 0.94), indicating that these measures capture highly concordant aspects of single-cell methylation disorder as well, whereas epiCHAOS showed moderately high correlations with PIM (r = 0.48) and SE (r = 0.61). These single-cell data support the conclusion that increased methylation entropy at PRC2-associated regions is a cell-intrinsic feature of cancer cells and is not solely attributable to differences in cellular composition within bulk tumor samples. Collectively, these analyses indicate that the genomic regions exhibiting the highest DNA methylation entropy across normal, premalignant, and malignant tissues and other aspects of the normal-to-cancer continuum are non-randomly associated with PRC2-regulated chromatin.

**Figure 3.**
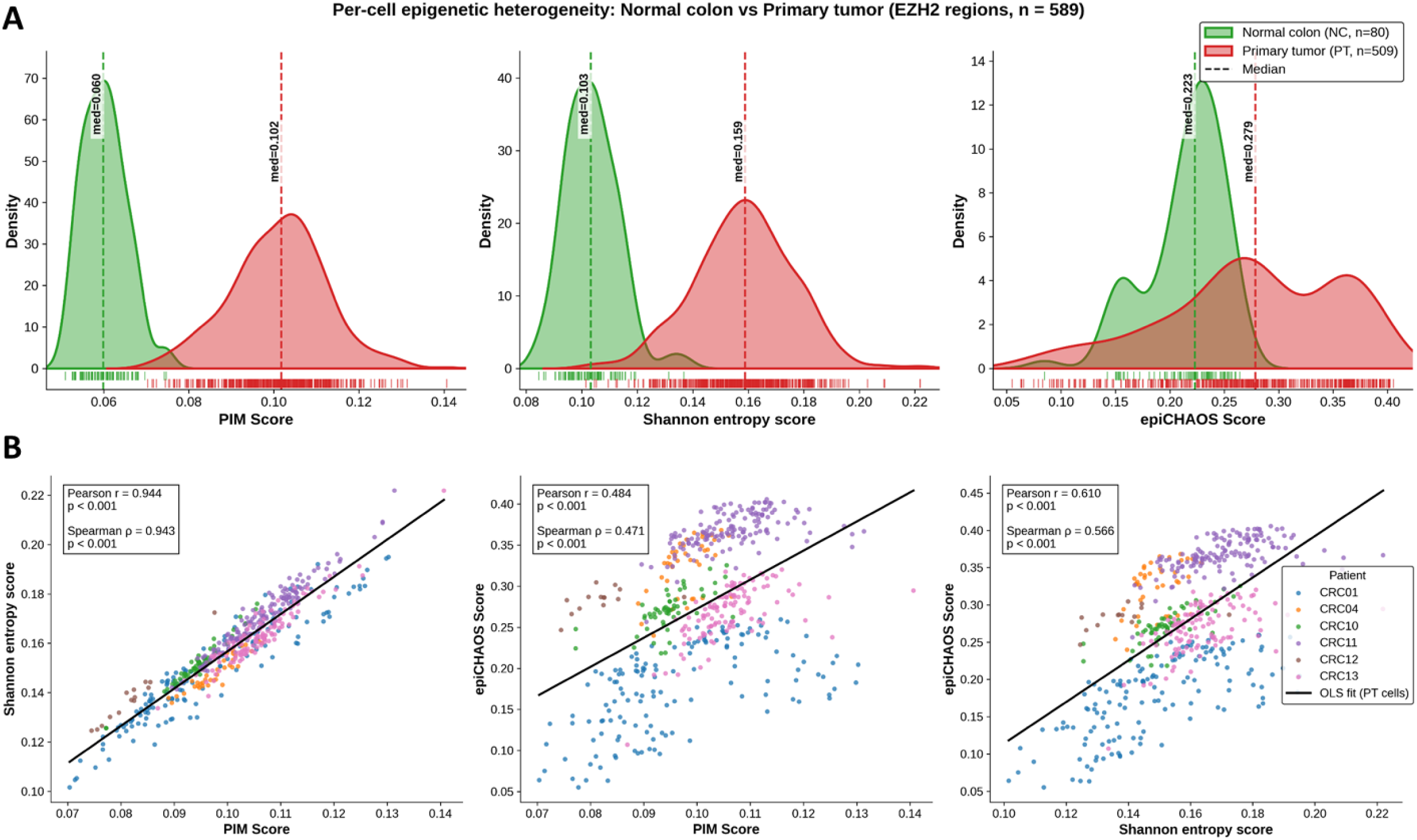
Single-cell DNA methylation disorder within EZH2-binding regions in normal colon and primary colorectal tumor cells. A) Density distributions of PIM, Shannon entropy, and epiCHAOS scores calculated per cell within EZH2-binding regions for normal colon cells (NC, n = 80) and primary tumor cells (PT, n = 509). Dashed vertical lines indicate group medians, and rug plots show individual cell-level values. B) Pairwise correlations among PIM, Shannon entropy, and epiCHAOS scores in primary tumor cells. Points are colored by patient (CRC01, CRC04, CRC10, CRC11, CRC12, and CRC13), and black lines indicate ordinary least-squares fits. Pearson and Spearman correlation coefficients and corresponding p values are shown within each panel.

### EZH2-binding intensity defines distinct DNA methylation trajectories across healthy, normal-adjacent, precancerous, and cancerous tissues

We next examined how DNA methylation levels and inter-sample variability change across EZH2-bound regions stratified by increasing EZH2 binding intensity (**Supplementary Figure 14**). EZH2 ChIP-seq peaks derived from human embryonic stem cells were used as a common proxy across cancer types. In the majority of cancers, mean DNA methylation increased progressively with increasing EZH2-binding rank, particularly in tumors, while substantial inter-sample methylation dispersion was observed across the full range of EZH2 occupancy. This pattern was substantially weaker or absent in matched normal tissues (**Figure 4**; **Supplementary Figure 15**). The same relationship was reproduced in the independent validation cohorts (**Figure 5**; **Supplementary Figure 16**). Notably, precancerous tissues closely recapitulated the methylation profiles observed in matched cancers, both with respect to sustained inter-sample variability and the progressive increase in DNA methylation with EZH2 binding intensity. In contrast, healthy tissues and normal-adjacent samples, including samples from different individuals and collected at varying distances from the tumor, showed highly overlapping methylation profiles that were clearly distinct from those observed in precancerous and cancerous tissues. Consistent with this PRC2-specific pattern, genome-wide analyses showed a comparable increase in DNA methylation dispersion when either tumors or precancerous lesions were compared with their respective normal tissues (**Supplementary Figure 17**). These findings indicate that the PRC2- and genome-wide associated methylation state characteristic of cancer is already established at the precancerous stage, consistent with early epigenetic reprogramming during tumorigenesis.

**Figure 4.**
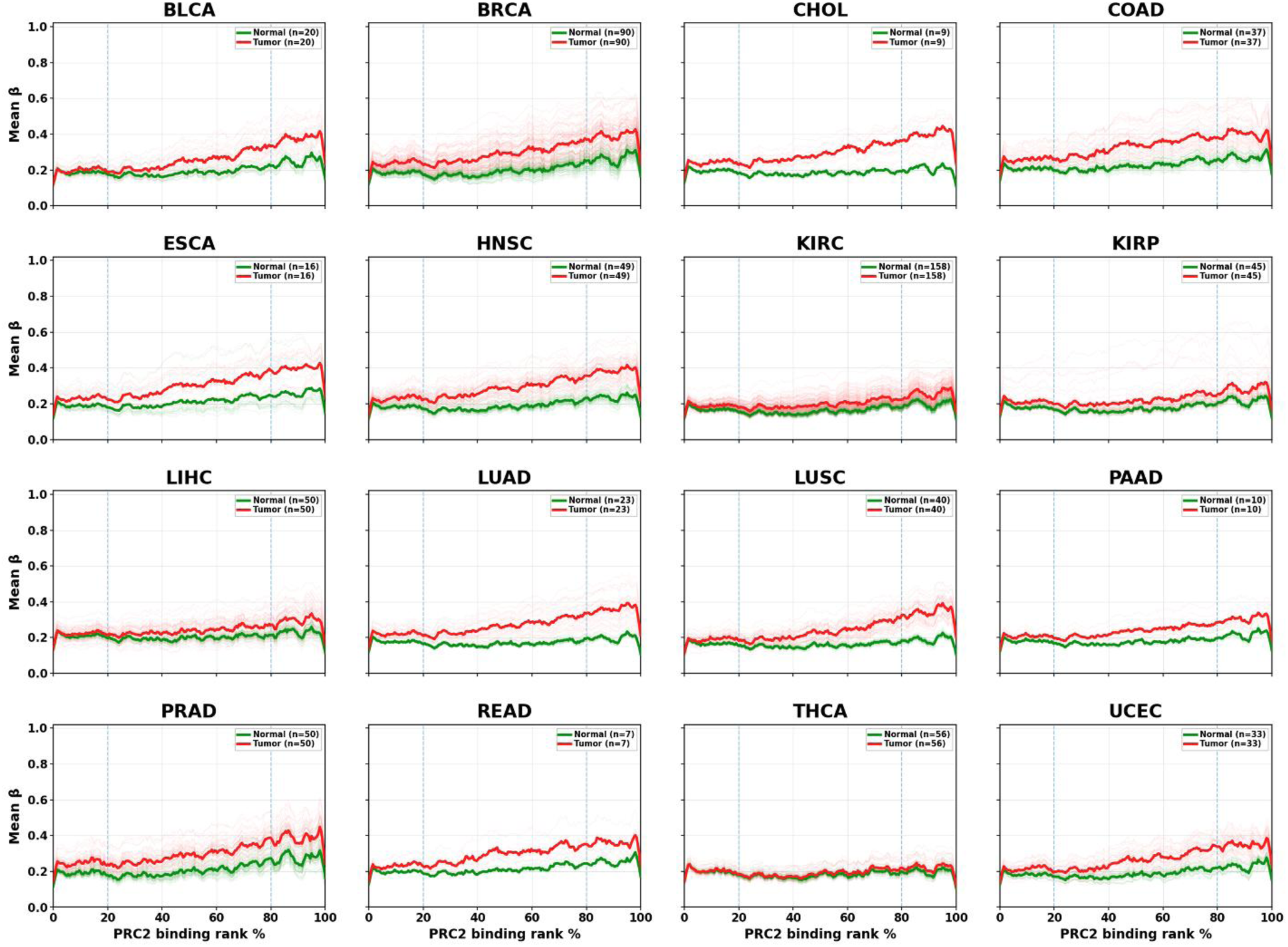
DNA methylation levels across increasing PRC2-binding intensity in matched normal and tumor tissues. Mean DNA methylation (β value) is plotted across genomic regions ranked from low to high PRC2 binding for 16 cancer types. Normal tissues are shown in green and matched tumor tissues in red. Thin lines represent individual samples, whereas solid lines represent the group mean. The x-axis indicates PRC2-binding rank percentile, and dashed vertical lines mark the 20th and 80th percentiles used to delineate low-, intermediate-, and high-binding regions. Sample numbers for normal and tumor tissues are indicated within each cancer-specific panel.

**Figure 5.**
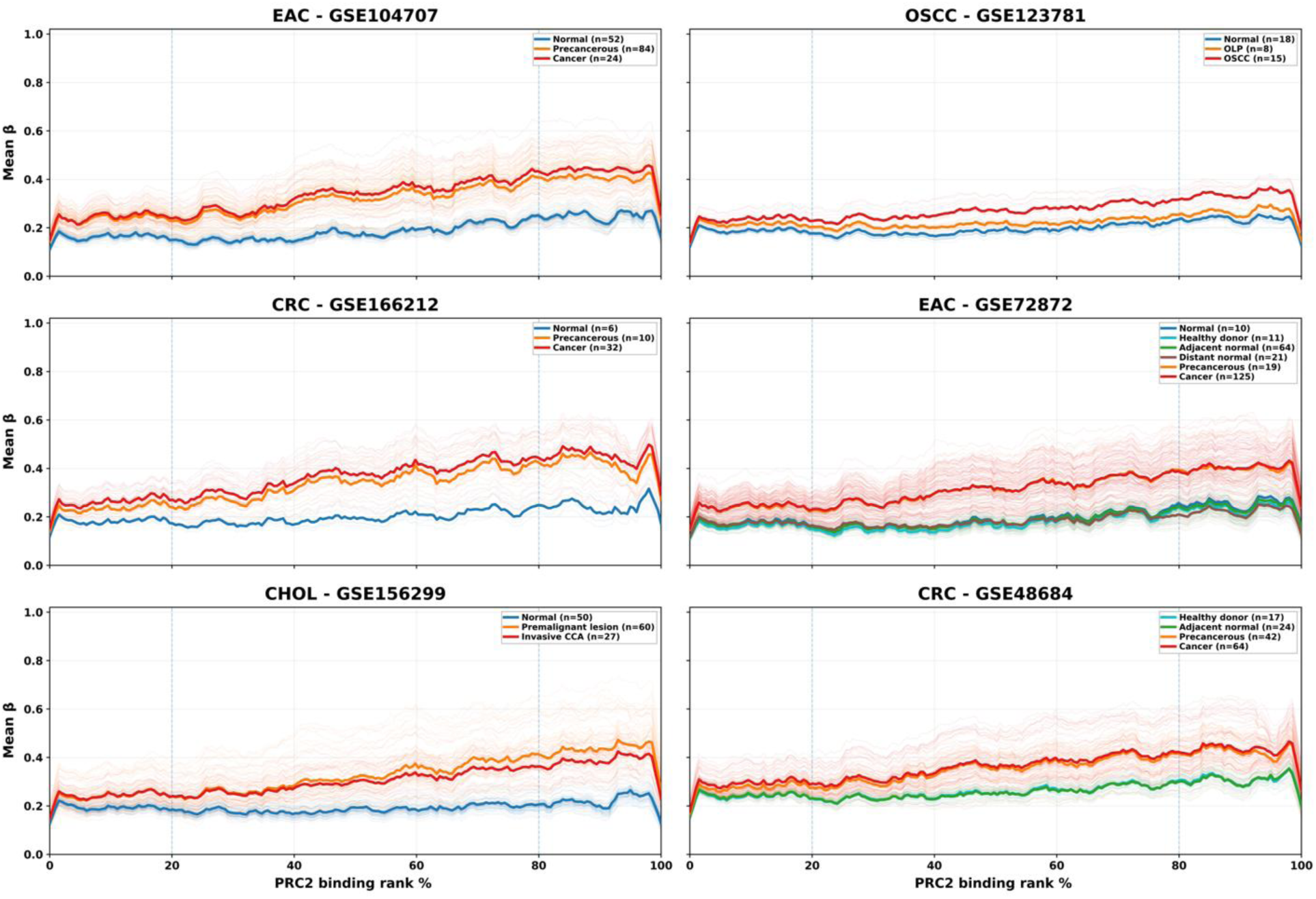
DNA methylation levels across increasing PRC2-binding intensity in independent validation cohorts spanning normal, precancerous, and cancerous tissues. Mean DNA methylation (β value) is plotted across genomic regions ranked from low to high PRC2 binding in six independent GEO datasets: esophageal adenocarcinoma (EAC; GSE104707 and GSE72872), oral squamous cell carcinoma (OSCC; GSE123781), colorectal cancer (CRC; GSE166212 and GSE48684), and cholangiocarcinoma (CHOL; GSE156299). Individual sample profiles are shown as thin lines and group means as solid lines. The x-axis indicates PRC2-binding rank percentile, and dashed vertical lines mark the 20th and 80th percentiles. Tissue categories and corresponding sample numbers are indicated within each panel.

### Low- and high-EZH2 regions occupy distinct regulatory chromatin environments

To determine whether methylated regions with low EZH2 occupancy, which retain methylation levels similar to normal tissue, differ in regulatory context from highly occupied EZH2 regions, which show increased DNA methylation, we compared their chromatin and regulatory annotations with LOLA (56) against publicly available chromatin and regulatory annotations from H1 human embryonic stem cells from Roadmap Epigenomics and Cistrome (57–59). H1 hESCs were used as a common reference because of their well-characterized chromatin architecture and relevance to developmental and cell identity.

Low-EZH2 regions were preferentially associated with transcription-related factors, including RNA polymerase and TAF7, and with chromatin features characteristic of more permissive regulatory states, including histone acetylation, H2A.Z, H3K4me3, and chromatin states annotated as weak, poised, and, to a lesser extent, active promoters. In contrast, high-EZH2 regions were enriched for H3K4me1, H4K20me1, H3K27me3 and, notably, H3K36me3, and were preferentially associated with poised promoters, repressed chromatin, and repetitive chromatin states (**Figure 6**; **Supplementary Figure 18**). Nearly identical enrichment patterns were observed in the validation cohort (**Supplementary Figures 19 and 20**).

**Figure 6.**
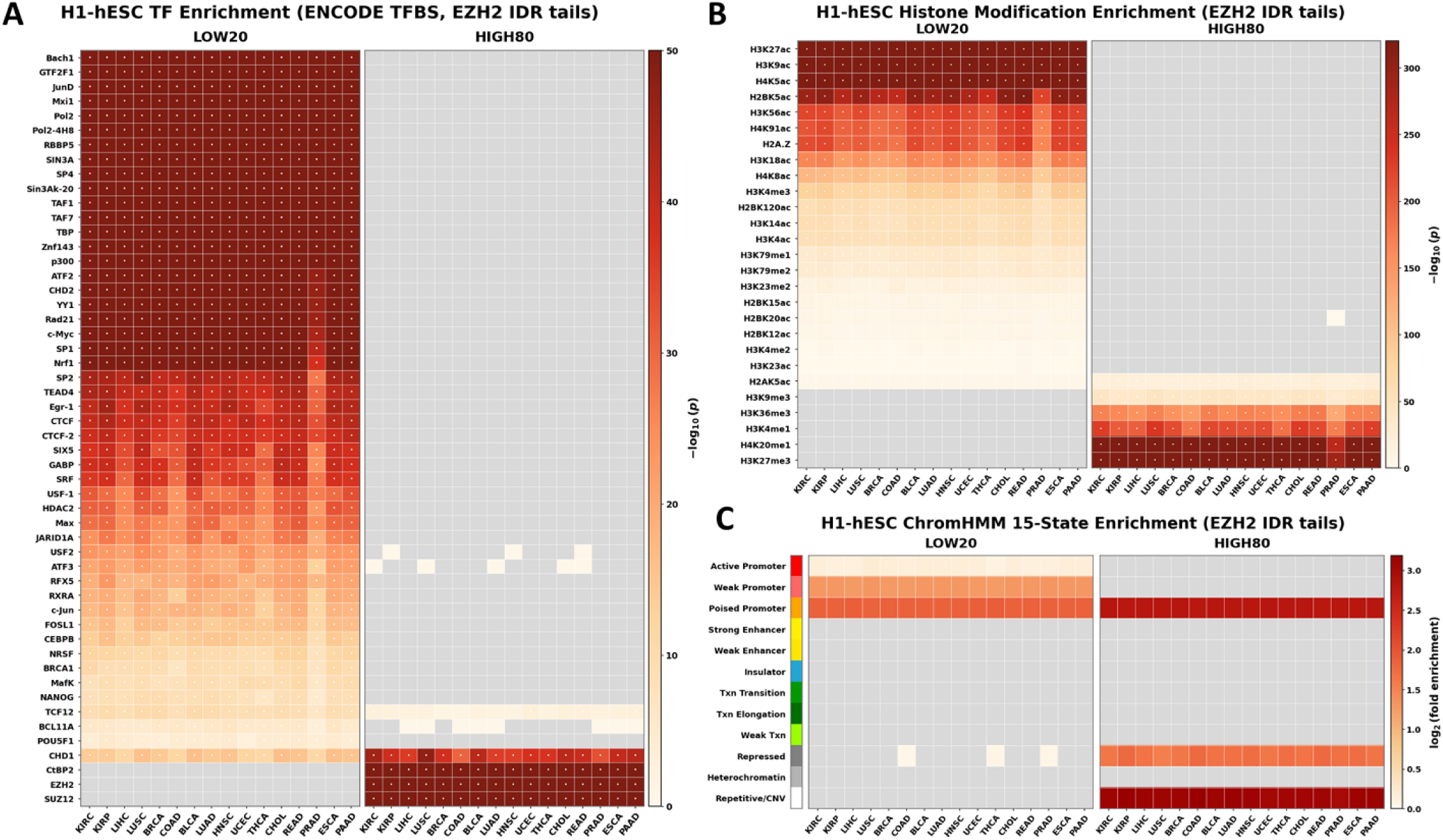
Regulatory and chromatin-state enrichment of low- and high-EZH2-binding regions. Genomic regions were stratified according to EZH2-binding intensity into the lowest 20% (LOW20) and highest 20% (HIGH80) and analyzed across the indicated cancer types using H1 human embryonic stem cell (H1-hESC) annotations. A) Enrichment of ENCODE transcription factor binding sites within LOW20 and HIGH80 EZH2-binding regions. Color intensity represents −log10(p) of the enrichment. B) Enrichment of histone modification annotations within LOW20 and HIGH80 EZH2-binding regions. Color intensity represents −log10(p) of the enrichment. C) Enrichment of H1-hESC ChromHMM 15-state chromatin annotations within LOW20 and HIGH80 EZH2-binding regions. Color intensity represents log2 fold enrichment. Gray cells indicate categories without enrichment values in the corresponding analysis.

Consistent with these annotations, high-EZH2 methylation regions exhibited lower chromatin accessibility by ATAC-seq than low-EZH2 regions and were associated with lower gene expression in both normal and tumor tissues (**Supplementary Figures 21 and 22**), indicating that EZH2 occupancy therefore defines a progressively less accessible and transcriptionally repressed chromatin state that may predispose these regions to aberrant DNA methylation during the transition from normal to precancerous and malignant tissue.

### Elevated methylation disorder at PRC2-associated regions is associated with aggressive tumor features and adverse survival

We next examined whether DNA methylation disorder within PRC2-associated regions was associated with clinical features of tumor aggressiveness. Higher PIM and Shannon entropy at PRC2 regions were positively associated with lymph node involvement at presentation and the occurrence of new neoplasm events following initial therapy (**Figure 7A and B**). Both metrics also increased with the Ragnum hypoxia score, associating PRC2-localized methylation disorder to a more hypoxic tumor state (**Figure 7C**).

**Figure 7.**
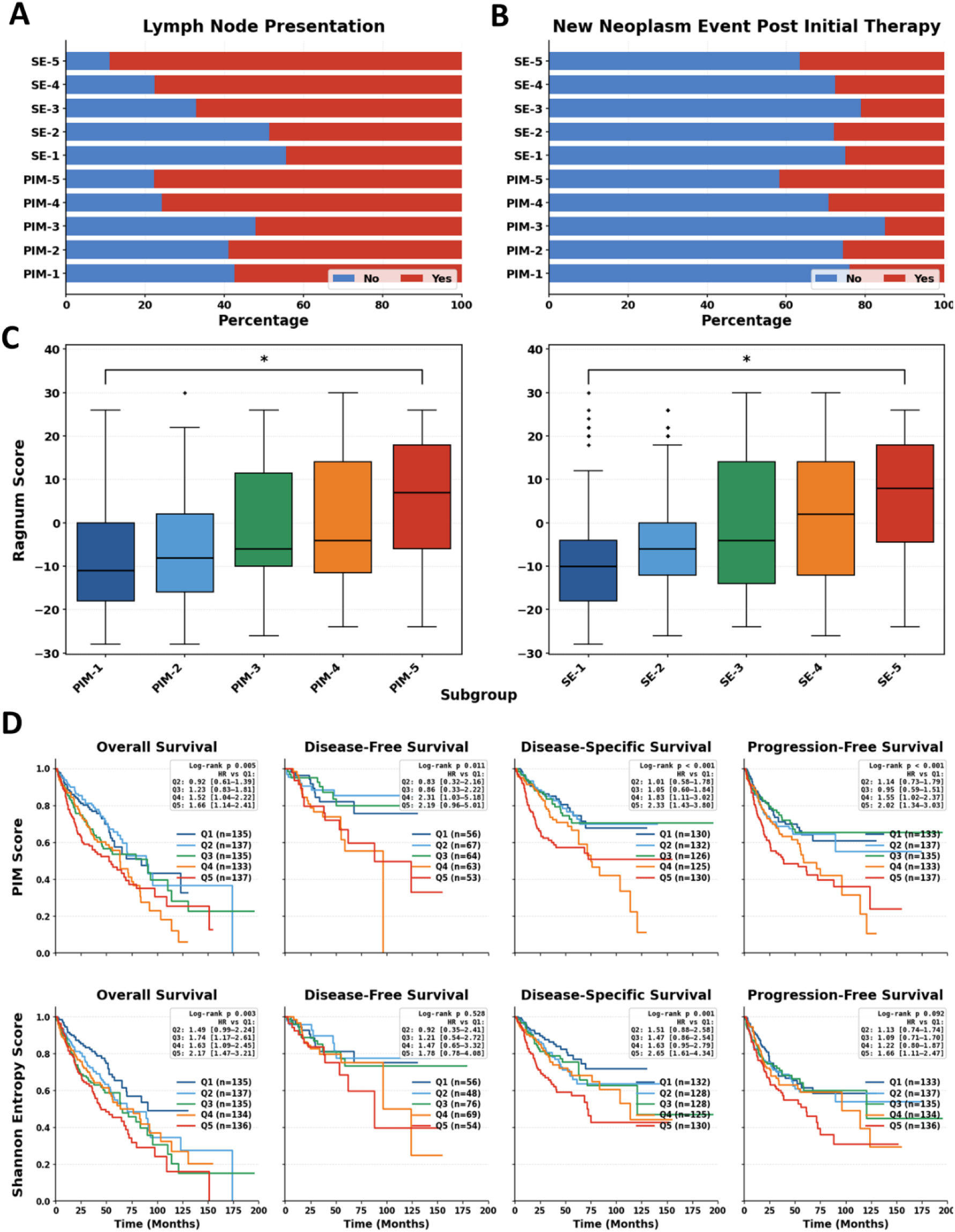
Clinical characteristics, hypoxia scores, and survival across PRC2-region DNA methylation disorder groups. A) Percentage of samples with or without lymph node involvement across quintiles of PIM and Shannon entropy (SE), with groups ordered from lowest (1) to highest (5) methylation disorder. B) Percentage of samples with or without a new neoplasm event after initial therapy across PIM and SE quintiles. C) Ragnum hypoxia scores across PIM and SE quintiles. Boxplots show the median, interquartile range, whiskers, and outliers; statistical significance is indicated above the corresponding comparisons (p < 0.05). D) Kaplan–Meier curves for overall survival, disease-free survival, disease-specific survival, and progression-free survival across PIM and SE quintiles. Insets show log-rank p values and hazard ratios with 95% confidence intervals relative to the lowest-disorder quintile (Q1).

We further assessed the relationship between PRC2-region methylation disorder and clinical outcome by stratifying tumors into quintiles according to PIM or SE. Increasing methylation entropy at EZH2 regions was enriched for deadlier cancer types (**Supplementary Figures 11C and F, and 23**). In line with that, increasing PIM was associated with significantly worse overall (log-rank p = 0.005), disease-free (p = 0.011), disease-specific (p < 0.001), and progression-free survival (p < 0.001) (**Figure 7D**). Compared with the lowest PIM quintile, the highest quintile showed increased hazards for overall survival (HR = 1.66, 95% CI 1.14–2.41), disease-free survival (HR = 2.19, 0.96–5.01), disease-specific survival (HR = 2.33, 1.43–3.80), and progression-free survival (HR = 2.02, 1.34–3.03). SE showed a similar direction of association, with significant separation for overall survival (p = 0.003; Q5 HR = 2.17, 1.47–3.21) and disease-specific survival (p = 0.001; Q5 HR = 2.65, 1.61–4.34). Progression-free survival showed a weaker overall trend (p = 0.092), although the highest SE quintile remained associated with increased risk (HR = 1.66, 1.11–2.47), whereas disease-free survival did not differ significantly across SE quintiles (p = 0.528). Overall, elevated DNA methylation disorder at PRC2-associated regions marks deadlier tumors with more aggressive clinical characteristics and adverse survival outcomes.

## DISCUSSION

In this study, we identify PRC2-associated chromatin as a major genomic compartment amenable to DNA methylation entropy across human tissues, especially cancers. Commencing with an unbiased analysis of transcription factor binding sites within regions of high methylation entropy, we consistently identified PRC2 components among the most enriched regulators in both normal and tumor tissues, with the same pattern reproduced in independent datasets. Stratification by EZH2 binding intensity in hESCs, which is the main enzymatic component of PRC2, further revealed a characteristic cancer-associated methylation profile with mean DNA methylation increasing progressively with EZH2 binding intensity, whereas inter-sample dispersion remained elevated across the binding range. This relationship was substantially weaker in normal tissues. Notably, precancerous lesions closely resembled malignant tissues in both methylation and dispersion levels across EZH2-binding strata and genome-wide, while healthy and tumor-adjacent tissues showed overlapping profiles that were distinct from the precancerous and malignant states. Regulatory annotation further showed that methylation in high-EZH2 regions preferentially occupied canonical PRC2 targets, such as poised and repressed chromatin, exhibited lower accessibility and lower transcriptional activity, whereas low-EZH2 regions were more frequently associated with permissive or poised-promoter and acetylated chromatin states. Single-cell colorectal cancer methylomes confirmed increased methylation entropy in PRC2 regions in malignant cells, arguing that the observed signal is likely not attributable to cellular admixture in bulk tumors. Finally, increased PIM and Shannon entropy within PRC2-associated regions were accompanied by more aggressive clinical features such as higher hypoxia scores, lymph-node involvement, new neoplasm events, and less favorable survival outcomes. Together, these findings indicate that PRC2-associated regulatory regions are hotspots for DNA methylation entropy.

PRC2-regulated loci provide a mechanistically relevant context for the observed methylation phenotype. PRC2 deposits H3K27me3 at developmental and lineage-regulatory regions, where it contributes to maintenance of transcriptional repression and cell-state identity. Many of these loci are poised or bivalent and therefore depend on tightly regulated interactions between repressive and activating chromatin marks. In our data, regions with higher EZH2 occupancy showed lower accessibility and transcriptional output, but in tumors also exhibited greater methylation gain and variability. This suggests that DNA methylation dysregulation preferentially affects a subset of PRC2-associated regulatory regions during transformation. Whether methylation gain at these loci is accompanied by loss of PRC2 occupancy or occurs while PRC2 remains bound cannot be resolved from the present datasets. Joint single-cell profiling of DNA methylation and PRC2/H3K27me3 occupancy will be required to distinguish these possibilities and to define the temporal and structural order of these epigenetic changes.

Interestingly, across validation cohorts, precancerous lesions already showed PRC2-associated methylation profiles resembling those of tumors, whereas healthy and tumor-adjacent tissues remained more similar to one another and were clearly distinct from the trends observed in precancerous and cancerous samples. This is consistent with substantial methylation remodeling at PRC2-associated and genome-wide regions occurring before invasive disease, although the cross-sectional design does not establish temporal order or causality.

Higher PRC2-region methylation disorder was associated with multiple clinically aggressive features such as lymph node involvement, post-therapy recurrence, hypoxia-associated transcriptional programs, and worse survival. These associations likely reflect coupled genomic and epigenomic instability rather than direct causal relationships. As in our previous analysis of global methylation entropy (26), higher PRC2-region PIM and Shannon entropy were associated with poorer survival, consistent with more aggressive tumor biology. However, these measures should not yet be considered independent prognostic biomarkers, as tumor type, stage, and other clinical covariates may confound the pan-cancer associations.

Several limitations should be considered. Most analyses rely on bulk Illumina 450K methylation data, which provide incomplete representation of repetitive elements, distal regulatory regions, and large heterochromatic domains. Although the single-cell colorectal dataset supports a cancer-cell-intrinsic component of the observed methylation disorder, validation across additional malignancies will be important. In addition, EZH2 and other chromatin-factor binding sites derived from H1 embryonic stem cells were used as a standardized reference across tumor types. This approach enables comparison of a common set of Polycomb-associated loci but does not capture tissue- or tumor-specific PRC2 occupancy, and it should be interpreted as a reference epigenome rather than direct measurements of occupancy in each tumor. Multi-matched methylome, EZH2/H3K27me3 profiling, chromatin accessibility, and transcriptomic data from the same tumors would provide a more direct assessment of these relationships.

Finally, the observational design of our study precludes causal inference. Functional and perturbational experiments will be required to establish whether altered PRC2 activity facilitates methylation disorder, whether DNA methylation changes modify PRC2 occupancy or function, or whether both reflect a broader loss of epigenetic regulatory fidelity.

Overall, our data identify PRC2-associated chromatin as a reproducible hotspot of DNA methylation gain and variability across normal, precancerous and malignant states. The findings suggest that the transition to cancer involves a pronounced reorganization of methylation at these developmentally regulated regions, with a cancer-like state already detectable in precancerous tissues. The positive correlation of this phenotype with hypoxia, aggressive clinical features, and adverse outcomes further indicates that methylation entropy in PRC2 regions captures a biologically relevant dimension of tumor epigenomic dysregulation. Whether this state contributes functionally to loss of lineage fidelity and increased cellular plasticity, or primarily reflects broader disruption of chromatin-state maintenance, remains an important question for future mechanistic studies.

## METHODS

### Datasets used in the study

Data were obtained from the cBioPortal for Cancer Genomics (https://www.cbioportal.org), the Genomic Data Commons (GDC) (https://portal.gdc.cancer.gov/), and the Gene Expression Omnibus (GEO).

As a discovery cohort, we analyzed paired Illumina HumanMethylation450 data from TCGA across 16 cancer types, comprising 693 matched tumor–adjacent normal pairs (LUSC, n = 40; BLCA, n = 20; KIRP, n = 45; UCEC, n = 33; BRCA, n = 90; READ, n = 7; ESCA, n = 16; LUAD, n = 23; PAAD, n = 10; HNSC, n = 49; CHOL, n = 9; THCA, n = 56; LIHC, n = 50; KIRC, n = 158; PRAD, n = 50; COAD, n = 37).

External validation comprised 18 independent GEO DNA methylation cohorts totaling 3,110 samples: GSE69914 (36) (breast cancer, n = 407; normal, n = 57; adjacent normal, n = 42; cancer, n = 308), GSE225845 (37) (breast cancer, n = 595; healthy donor, n = 231; adjacent normal, n = 140; cancer, n = 224), GSE287331 (38) (breast cancer, n = 446; healthy donor, n = 182; adjacent normal, n = 60; contralateral normal, n = 68; opposite-quadrant normal, n = 67; cancer, n = 69), GSE133985 (39) (breast cancer, n = 25; adjacent normal, n = 8; contralateral normal, n = 9; cancer, n = 8), GSE66313 (40) (breast DCIS, n = 55; adjacent normal, n = 15; DCIS, n = 40), GSE199057 (41) (colorectal cancer, n = 221; healthy donor, n = 68; normal tissue from the CRC cohort, n = 77; cancer, n = 76), GSE48684 (42) (colorectal cancer, n = 147; healthy donor, n = 17; adjacent normal, n = 24; adenoma, n = 42; cancer, n = 64), GSE166212 (43) (colorectal cancer, n = 48; normal, n = 6; adenoma, n = 10; carcinoma, n = 32), GSE56044 (44) (lung cancer, n = 136; adjacent normal, n = 12; carcinoma, n = 124), GSE108123 (45) (pre-invasive lung disease, n = 87; control, n = 33; regressive lesion, n = 18; progressive lesion, n = 36), GSE89852 (46) (hepatocellular carcinoma, n = 74; non-cancerous liver, n = 37; HCC, n = 37), GSE112047 (47) (prostate cancer, n = 47; adjacent normal, n = 16; cancer, n = 31), GSE116338 (48) (prostate cancer progression, n = 43; benign, n = 10; PIA, n = 7; HGPIN, n = 6; primary cancer, n = 14; metastatic cancer, n = 6), GSE104707 (49) (esophageal adenocarcinoma, n = 160; normal, n = 52; precancerous, n = 84; EAC, n = 24), GSE72872 (50) (esophageal adenocarcinoma, n = 250; normal, n = 10; healthy donor, n = 11; adjacent normal, n = 64; distant/stomach tissue, n = 21; precancerous, n = 19; EAC, n = 125), GSE103186 (51) (gastric premalignant disease, n = 191; normal, n = 61; intestinal metaplasia, n = 130), GSE156299 (52) (cholangiocarcinoma, n = 137; normal bile duct, n = 50; premalignant lesion, n = 60; invasive CCA, n = 27), and GSE123781 (53) (oral squamous cell carcinoma, n = 41; normal, n = 18; oral lichen planus, n = 8; OSCC, n = 15).

### DNA methylation profiles

Genome-wide DNA methylation profiles from TCGA were generated using the Illumina HumanMethylation450 (450K) BeadChip platform, which interrogates over 450,000 CpG sites spanning promoter regions, gene bodies, CpG islands, shores, shelves, and open-sea regions. Harmonized and quality-controlled Level 3 β-values, ranging from 0 (unmethylated) to 1 (fully methylated), were retrieved via the GDC application programming interface (API). Samples were matched by TCGA patient identifier to construct tumor–adjacent normal pairs, resulting in 693 matched pairs across 16 cancer types. Methylation matrices were assembled and processed on a high-performance computing cluster. Independent validation was performed using 18 publicly available DNA methylation datasets obtained primarily from the Gene Expression Omnibus (GEO), encompassing healthy tissue, tumor-adjacent normal tissue, normal tissue sampled at different distances from the tumor, precancerous lesions, invasive cancers, and progressive or regressive pre-invasive lesions. These datasets were generated using either the Illumina HumanMethylation450 BeadChip or the Illumina MethylationEPIC array. For 17 datasets, publisher-provided processed β-value matrices were used, whereas GSE225845 was processed from raw IDAT files. No additional cross-study normalization of β-values was performed, and each validation cohort was analyzed independently according to its corresponding sample annotations and biological groups.

For both TCGA and validation datasets, CpG probes with missing measurements and probes mapping to sex chromosomes were excluded where appropriate. Probe annotations were harmonized to the hg19 genome assembly to ensure consistent genomic coordinates across the 450K and EPIC platforms and compatibility with genomic annotation and ChIP-seq datasets. Probes with conflicting genomic coordinates were excluded, duplicate probe identifiers were resolved, and, when probes from both platforms mapped to the same genomic position, the EPIC annotation was preferentially retained. The resulting standardized methylation matrices were used for downstream analyses.

Subsequent analyses were performed on EZH2-associated genomic regions as a functionally relevant component of the PRC2 complex. For this purpose, publicly available EZH2 ChIP-seq peaks (ENCODE accession ENCSR000ASY) from the Encyclopedia of DNA Elements (ENCODE) consortium were used (60). The peaks were downloaded in narrowPeak format using hg19/GRCh37 genomic coordinates. The dataset contained 18,589 IDR-ranked peaks (Irreproducible Discovery Rate), enabling the selection of highly reproducible ChIP-seq signals detected across biological replicates.

Clinicogenomic data were retrieved from TCGA clinical metadata deposited in cBioPortal. Chromatin accessibility data were obtained from the GDC portal. RNA-seq data were downloaded from the GDC portal.

### Methylation entropy metrics: PIM and Shannon entropy

PRC2-associated intermediate methylation was quantified using the Proportion of Intermediate Methylation (PIM). Illumina HumanMethylation450K β-values were used as provided from the preprocessed β-value matrix. For each sample, CpG sites were classified as intermediately methylated if their β-values fell within the range 0.2 ≤ β < 0.6, and PIM was calculated as the number of CpGs in EZH2-binding regions in this intermediate range divided by the total number of retained CpGs for that sample in EZH2-binding regions. Tumor–normal differences in PIM were visualized using overlaid histograms and cancer-stratified boxplots, and statistical significance was assessed using two-sided Mann–Whitney U tests for unpaired comparisons and paired Wilcoxon signed-rank tests for matched samples.

PRC2-associated DNA methylation disorder was quantified at the sample level using Shannon entropy (61) After standard preprocessing of Illumina HumanMethylation450K arrays, β-values were treated as Bernoulli parameters for each CpG site and sample. To avoid numerical artefacts, β-values were constrained to the interval [*ε*, 1 − *ε*] (*ε* = 1 × 10⁻⁶). CpG-level entropy was computed as

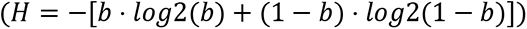

and a single per-sample entropy score was calculated as the mean entropy across all retained CpGs.

CpG positions were intersected with ChIP-seq peak regions for all 49 transcription factors based on genomic overlap. CpGs falling within a ChIP-seq peak for a given transcription factor were classified as associated with that factor. Because probe coverage differed between the Illumina HumanMethylation450K and EPIC platforms, each validation cohort was analyzed using the transcription factor-associated CpGs available within that specific dataset, rather than restricting the analysis to probes shared across all platforms and studies. This procedure was applied consistently across all 49 transcription factors, including EZH2. Random control sets were generated for the CHD1- and CTCF-associated regions, containing the same number of CpG sites as the corresponding target set. Control CpG sites were selected from outside the CHD1/CTCF regions while preserving the same chromosome-wise distribution of CpGs as in the target set.

### Single-cell DNA methylation analysis

To validate PRC2-associated DNA methylation heterogeneity at the single-cell level, we analyzed publicly available single-cell whole-genome bisulfite sequencing (scWGBS) data from colorectal cancer generated using scTrioSeq2 (GSE97693) (54). The dataset comprised 589 cells from six patients, including 80 normal epithelial cells and 509 primary tumor cells. CpG-context methylation calls were extracted from single-cell methylation files and aggregated into fixed 1-kb genomic windows across autosomes and chromosome X after coverage-based filtering. Only windows covered by at least three reads per cell and present in at least five tumor and five normal cells per patient were retained for downstream analyses.

PRC2-associated epigenetic heterogeneity was quantified using three complementary metrics: epiCHAOS, the proportion of intermediately methylated regions (PIM), and Shannon entropy. ЕpiCHAOS was calculated according to the original publication (55) by binarizing methylation states within 1-kb windows using a threshold of 0.5 (greater than or equal to 0.5 were treated as methylated and windows with average methylation below 0.5 were treated as unmethylated), computing pairwise Jaccard similarities between cells, correcting for expected similarity and methylation sparsity, and transforming the resulting residuals into a heterogeneity score in which higher values indicate greater epigenetic disorder. All analyses were conducted in PRC2-associated regions.

### DNA methylation in PRC2 regions

For visualization DNA methylation in PRC2-associated regions, EZH2 ChIP-seq peaks were ranked from low to high signal intensity and divided into 200 consecutive bins. CpG probes overlapping the ranked peaks were assigned to the corresponding bins, and mean β-values were calculated per sample and summarized separately for tumor and matched normal tissues within each cancer type. For visualization, the resulting curves were smoothed using a seven-bin moving average to reduce local fluctuations. Smoothing was used only for graphical representation and did not affect the CpG-level data used for statistical analyses.

For complementary distribution-based comparisons, the ranked EZH2-associated CpGs were divided into ten deciles (D1–D10), representing progressively increasing EZH2 binding intensity. Methylation values were summarized per sample within each decile and visualized using boxplots for tumor and matched normal tissues within each cancer type. Differences between tumor and normal samples were evaluated separately for each decile.To further characterize genomic features associated with the extremes of EZH2 occupancy, EZH2-associated regions were stratified according to ChIP-seq signal intensity. Regions within the lowest 20% of the EZH2 signal distribution were designated as LOW20, whereas regions at or above the 80th percentile, corresponding to the highest 20%, were designated as HIGH80. These two groups were subsequently compared across complementary genomic, epigenomic, chromatin-accessibility, and transcriptomic features to characterize functional differences associated with low versus high EZH2 occupancy.

### Genomic and chromatin state annotations

Regulatory enrichment analyses were performed using LOLA (Locus Overlap Analysis) (56). Enrichment was evaluated against publicly available ChIP-seq peak annotations from the H1 human embryonic stem cell (H1-hESC) context and tissue, including ENCODE transcription factor binding sites and Roadmap Epigenomics histone modification datasets, accessed via the LOLA RegionDB (hg19). The same was done for chromatin-state annotation using the hg19 ChromHMM segmentations (62) taken from H1-hESC. Chromatin-state distributions were summarized relative to a 450K-matched background to assess enrichment or depletion of unstable regions across major chromatin states. Furthermore, tissue specificity was evaluated by repeating the enrichment and annotation analyses using chromatin segmentation and histone modification datasets from primary cell types representative of the corresponding tumor tissue of origin.

Overlap enrichment was quantified using odds ratios, q-values, and p-values, with multiple-testing correction applied where appropriate, and enrichment signals were summarized across cancer types for downstream analyses. The enrichment metric was selected based on the clarity and interpretability of the resulting visualizations.

Genomic annotation of stable and unstable DNA methylation variability regions was performed using pyBEDtools (63). BED-formatted CpG-centered regions representing stable and unstable sets were annotated against UCSC hg19 gene annotations, while CpG-island context was assigned using Illumina HumanMethylation450K probe annotations; the full 450K probe set was used as the genomic background.

### ATAC-seq Analysis of Chromatin Accessibility in EZH2-Associated Regions

Chromatin accessibility of EZH2-associated regions was analyzed to assess whether LOW20 and HIGH80 regions differ in local chromatin accessibility. Previously processed ATAC-seq insertion-signal tracks in BigWig format were used for 14 cancer types: BLCA, BRCA, CHOL, COAD, ESCA, HNSC, KIRC, KIRP, LIHC, LUAD, LUSC, PRAD, THCA, and UCEC. Corresponding ATAC-seq data were not available for PAAD and READ. Only autosomal chromosomes (chr1–chr22) were included in the analysis, whereas sex chromosomes were excluded (64). The data were processed using pyBigWig v0.3.24. For each LOW20 and HIGH80 region, a ±2 kb window around the center of the region was defined and divided into 200 intervals of 20 bp each. The mean ATAC-seq insertion signal was calculated for each interval, yielding centered chromatin-accessibility profiles for each cancer type. Chromatin accessibility was additionally quantified using the mean ATAC-seq signal in the central ±200 bp window around the center of the region. Differences in mean ATAC-seq signal between LOW20 and HIGH80 regions were tested separately for each of the 14 cancer types using a two-sided Mann-Whitney U test. The magnitude and direction of the effect were expressed as the log₂ fold change (log₂FC) between the mean values of the two groups. The resulting p-values were corrected for multiple testing across cancer types using the Benjamini-Hochberg (BH) procedure. In addition to peak-centered chromatin-accessibility profiles within a ±2 kb window around the region center, ATAC-seq signal was also quantified for the same LOW20 and HIGH80 regions within the central ±200 bp window. This analysis was used for statistical comparison of local chromatin accessibility between the two groups. The mean ATAC-seq signal in the central window was compared between LOW20 and HIGH80 regions using a two-sided Mann-Whitney U test, separately for each cancer type

### RNA-seq analysis of EZH2-Associated Regions

The transcriptional activity of genes associated with EZH2-associated regions was analyzed using RNA-seq data to assess whether differences between LOW20 and HIGH80 regions were reflected at the level of gene expression. RNA-seq STAR-augmented gene-count files containing gene-expression values expressed as transcripts per million (TPM) were used for this purpose. The analysis was performed on 370 tumor and paired normal samples (65).

LOW20 and HIGH80 EZH2-associated regions were supplied as BED files and annotated relative to promoter windows of ±2 kb around the transcription start site (TSS). Promoter coordinates were obtained from the UCSC hg19 knownGene annotation, and overlap with LOW20 and HIGH80 regions was used to define the corresponding gene sets. Genes whose promoters overlapped at least one region from the corresponding group were included in the LOW20 or HIGH80 gene set. For integration with RNA-seq data, genomic annotations and gene identifiers were matched to the corresponding RNA-seq gene-expression values. These steps were performed in R using the Bioconductor packages rtracklayer, GenomicRanges, TxDb.Hsapiens.UCSC.hg19.knownGene, org.Hs.eg.db, and AnnotationDbi (66–69).

### Analyses of clinical features

Clinical characterization of methylation heterogeneity was performed to assess whether epigenetic heterogeneity is associated with clinical and molecular characteristics of tumor samples. Two independent measures of epigenetic heterogeneity were used for each sample, PIM and SE, and the values were stratified into five quintile groups with approximately equal numbers of samples (Q1–Q5), with Q1 representing samples with the lowest and Q5 samples with the highest epigenetic heterogeneity.

Clinical and molecular data were obtained through the cBioPortal REST API. Lymph-node involvement status, occurrence of a new neoplastic event after initial treatment, and the Ragnum tumor hypoxia score were also analyzed. Differences between the Q1 and Q5 groups were tested using a two-sided Mann-Whitney U test.

The association between epigenetic heterogeneity and clinical outcomes was additionally evaluated using Kaplan-Meier survival analysis. Four clinical outcomes were analyzed: overall survival (OS), disease-free survival (DFS), progression-free survival (PFS), and disease-specific survival (DSS). For OS, the event was defined as death from any cause; for DFS, recurrence or progression; for PFS, disease progression; and for DSS, tumor-related death. Differences in survival between groups were assessed using the log-rank test (70). In addition, a univariable Cox proportional hazards model was applied to quantitatively assess differences in risk between quintile groups. Hazard ratios (HRs) and 95% confidence intervals (CIs) were calculated for groups Q2–Q5, with Q1 used as the reference group.

All statistical analyses were performed in Python v3.11. The scipy.stats package was used for nonparametric statistical tests, lifelines v0.27+ for survival analysis, and matplotlib v3.7+ for visualization and generation of graphical outputs.

### Statistics and visualizations

Statistical analyses included two-sided Wilcoxon signed-rank tests for paired comparisons, Mann– Whitney U tests for independent groups, DiffVar/missMethyl package using a limma-based empirical Bayes framework, for differential methylation variability analysis, χ² and Fisher’s exact tests for categorical and enrichment analyses, Pearson and Spearman correlations, bootstrap confidence intervals, BH-FDR and Bonferroni multiple-testing correction, and Kaplan–Meier, log-rank, and Cox proportional-hazards analyses for survival. Effect sizes were reported using Cliff’s δ, paired Cohen’s d, Cramér’s V, odds ratios, and log₂ fold-changes, as appropriate. Computational analyses were performed in Python 3.11 and R/Bioconductor using NumPy, SciPy, Matplotlib, lifelines, pyBEDTools, pyBigWig, missMethyl/DiffVar, ChIPseeker, GenomicRanges, GenomeInfoDb, rtracklayer, AnnotationDbi, LOLA. Genomic annotation used the hg19/GRCh37 reference and associated Bioconductor annotation databases, while clinical and molecular data were retrieved through the cBioPortal REST API.

## Declarations

### Ethics approval and consent to participate

Not applicable. All data are taken from publicly available datasets and cancer projects.

### Consent for publication

Not applicable. All data are taken from publicly available datasets and already reported in the literature.

### Availability of data and materials

All data are publicly available at the cBioportal repository, https://www.cbioportal.org/, and the Genomic Data Commons data portal https://portal.gdc.cancer.gov/. For scWGBS, we used the GSE97693 dataset. For external validation GSE69914, GSE225845, GSE287331, GSE133985, GSE66313, GSE199057, GSE48684, GSE166212, GSE56044, GSE108123, GSE89852, GSE112047, GSE116338, GSE104707, GSE72872, GSE103186, GSE156299 and GSE123781.

### Use of AI

ChatGPT-5.6 Sol and Grammarly were utilized for language editing and stylistic improvement. No AI tools were used for data analysis, data interpretation, or the generation of scientific conclusions. All content was reviewed and approved by the authors.

### Author contributions

GK conceived, designed, and led the study. MSM supervised aspects of the computational and statistical analyses. DjB collected, curated, processed, and analyzed the data with the help of GK. MGj added intellectual input to the study. GK wrote the initial draft of the manuscript. All authors contributed to data interpretation, critically revised the manuscript for important intellectual content, and approved the final version.

## REFERENCES

1. Timp W, Feinberg AP. Cancer as a dysregulated epigenome allowing cellular growth advantage at the expense of the host. Nat Rev Cancer. 2013 Jul;13(7):497–510. doi:10.1038/nrc3486 PubMed PMID: 23760024; PubMed Central PMCID: PMC4636434.

2. Easwaran H, Tsai HC, Baylin SB. Cancer epigenetics: tumor heterogeneity, plasticity of stem-like states, and drug resistance. Mol Cell. 2014 Jun 5;54(5):716–27. doi:10.1016/j.molcel.2014.05.015 PubMed PMID: 24905005; PubMed Central PMCID: PMC4103691.

3. Irizarry RA, Ladd-Acosta C, Wen B, Wu Z, Montano C, Onyango P, et al. The human colon cancer methylome shows similar hypo- and hypermethylation at conserved tissue-specific CpG island shores. Nat Genet. 2009 Feb;41(2):178–86. doi:10.1038/ng.298 PubMed PMID: 19151715; PubMed Central PMCID: PMC2729128.

4. Fernandez AF, Assenov Y, Martin-Subero JI, Balint B, Siebert R, Taniguchi H, et al. A DNA methylation fingerprint of 1628 human samples. Genome Res. 2012 Feb;22(2):407–19. doi:10.1101/gr.119867.110 PubMed PMID: 21613409; PubMed Central PMCID: PMC3266047.

5. Weisenberger DJ. Characterizing DNA methylation alterations from The Cancer Genome Atlas. J Clin Invest. 2014 Jan;124(1):17–23. doi:10.1172/JCI69740 PubMed PMID: 24382385; PubMed Central PMCID: PMC3871233.

6. Saghafinia S, Mina M, Riggi N, Hanahan D, Ciriello G. Pan-Cancer Landscape of Aberrant DNA Methylation across Human Tumors. Cell Rep. 2018 Oct 23;25(4):1066–1080.e8. doi:10.1016/j.celrep.2018.09.082 PubMed PMID: 30355485.

7. Hoadley KA, Yau C, Hinoue T, Wolf DM, Lazar AJ, Drill E, et al. Cell-of-Origin Patterns Dominate the Molecular Classification of 10,000 Tumors from 33 Types of Cancer. Cell. 2018 Apr 5;173(2):291–304.e6. doi:10.1016/j.cell.2018.03.022 PubMed PMID: 29625048; PubMed Central PMCID: PMC5957518.

8. Mazor T, Pankov A, Song JS, Costello JF. Intratumoral Heterogeneity of the Epigenome. Cancer Cell. 2016 Apr 11;29(4):440–51. doi:10.1016/j.ccell.2016.03.009 PubMed PMID: 27070699; PubMed Central PMCID: PMC4852161.

9. Guo M, Peng Y, Gao A, Du C, Herman JG. Epigenetic heterogeneity in cancer. Biomark Res. 2019;7:23. doi:10.1186/s40364-019-0174-y PubMed PMID: 31695915; PubMed Central PMCID: PMC6824025.

10. Witte T, Plass C, Gerhauser C. Pan-cancer patterns of DNA methylation. Genome Med. 2014;6(8):66. doi:10.1186/s13073-014-0066-6 PubMed PMID: 25473433; PubMed Central PMCID: PMC4254427.

11. Hanahan D. Hallmarks of cancer-Then and now, and beyond. Cell. 2026 Apr 16;189(8):2254–77. doi:10.1016/j.cell.2025.12.049 PubMed PMID: 41616779.

12. Hansen KD, Timp W, Bravo HC, Sabunciyan S, Langmead B, McDonald OG, et al. Increased methylation variation in epigenetic domains across cancer types. Nat Genet. 2011 Jun 26;43(8):768–75. doi:10.1038/ng.865 PubMed PMID: 21706001; PubMed Central PMCID: PMC3145050.

13. Pisanic TR, Athamanolap P, Wang TH. Defining, distinguishing and detecting the contribution of heterogeneous methylation to cancer heterogeneity. Semin Cell Dev Biol. 2017 Apr;64:5–17. doi:10.1016/j.semcdb.2016.08.030 PubMed PMID: 27582426; PubMed Central PMCID: PMC6339831.

14. Teschendorff AE. On epigenetic stochasticity, entropy and cancer risk. Philos Trans R Soc Lond B Biol Sci. 2024 Apr 22;379(1900):20230054. doi:10.1098/rstb.2023.0054 PubMed PMID: 38432318; PubMed Central PMCID: PMC10909509.

15. Aryee MJ, Liu W, Engelmann JC, Nuhn P, Gurel M, Haffner MC, et al. DNA methylation alterations exhibit intraindividual stability and interindividual heterogeneity in prostate cancer metastases. Sci Transl Med. 2013 Jan 23;5(169):169ra10. doi:10.1126/scitranslmed.3005211 PubMed PMID: 23345608; PubMed Central PMCID: PMC3577373.

16. Ma S, Pan X, Gan J, Guo X, He J, Hu H, et al. DNA methylation heterogeneity attributable to a complex tumor immune microenvironment prompts prognostic risk in glioma. Epigenetics. 2024 Dec;19(1):2318506. doi:10.1080/15592294.2024.2318506 PubMed PMID: 38439715; PubMed Central PMCID: PMC10936651.

17. Quek K, Li J, Estecio M, Zhang J, Fujimoto J, Roarty E, et al. DNA methylation intratumor heterogeneity in localized lung adenocarcinomas. Oncotarget. 2017 Mar 28;8(13):21994–2002. doi:10.18632/oncotarget.15777 PubMed PMID: 28423542; PubMed Central PMCID: PMC5400640.

18. Klughammer J, Kiesel B, Roetzer T, Fortelny N, Nemc A, Nenning KH, et al. The DNA methylation landscape of glioblastoma disease progression shows extensive heterogeneity in time and space. Nat Med. 2018 Oct;24(10):1611–24. doi:10.1038/s41591-018-0156-x PubMed PMID: 30150718; PubMed Central PMCID: PMC6181207.

19. Rossi SH, Dombrowe V, Godfrey L, Bucaciuc Mracica T, Pita S, Milne-Clark T, et al. Evidence of DNA methylation heterogeneity and epipolymorphism in kidney cancer tissue samples. Oncogene. 2025 Apr;44(15):1024–36. doi:10.1038/s41388-024-03270-3 PubMed PMID: 39824946; PubMed Central PMCID: PMC11976292.

20. Hua X, Zhao W, Pesatori AC, Consonni D, Caporaso NE, Zhang T, et al. Genetic and epigenetic intratumor heterogeneity impacts prognosis of lung adenocarcinoma. Nat Commun. 2020 May 18;11(1):2459. doi:10.1038/s41467-020-16295-5 PubMed PMID: 32424208; PubMed Central PMCID: PMC7235245.

21. Teng H, Xue M, Liang J, Wang X, Wang L, Wei W, et al. Inter- and intratumor DNA methylation heterogeneity associated with lymph node metastasis and prognosis of esophageal squamous cell carcinoma. Theranostics. 2020;10(7):3035–48. doi:10.7150/thno.42559 PubMed PMID: 32194853; PubMed Central PMCID: PMC7053185.

22. Gaiti F, Chaligne R, Gu H, Brand RM, Kothen-Hill S, Schulman RC, et al. Epigenetic evolution and lineage histories of chronic lymphocytic leukaemia. Nature. 2019 May;569(7757):576–80. doi:10.1038/s41586-019-1198-z PubMed PMID: 31092926; PubMed Central PMCID: PMC6533116.

23. Wang Y, Zhang Y, Huang Y, Chen C, Zhang X, Xing Y, et al. Intratumor heterogeneity of breast cancer detected by epialleles shows association with hypoxic microenvironment. Theranostics. 2021;11(9):4403–20. doi:10.7150/thno.53737 PubMed PMID: 33754068; PubMed Central PMCID: PMC7977462.

24. Monyak DL, Holloway ST, Gumbert GJ, Grimm LJ, Hwang ES, Marks JR, et al. Mapping the temporal landscape of breast cancer using epigenetic entropy. Commun Biol. 2025 Oct 16;8(1):1477. doi:10.1038/s42003-025-08867-2 PubMed PMID: 41102398; PubMed Central PMCID: PMC12533048.

25. McGranahan N, Swanton C. Biological and therapeutic impact of intratumor heterogeneity in cancer evolution. Cancer Cell. 2015 Jan 12;27(1):15–26. doi:10.1016/j.ccell.2014.12.001 PubMed PMID: 25584892.

26. Bukovec D, Gjorgjioski B, Misheva MS, Kungulovski G. DNA methylation variability defines a fundamental dimension of tumor epigenomes linked to genomic instability, tumor aggressiveness, and clinical outcomes. Epigenetics Chromatin. 2026 Sep 29. doi:10.1186/s13072-026-00704-y

27. Landan G, Cohen NM, Mukamel Z, Bar A, Molchadsky A, Brosh R, et al. Epigenetic polymorphism and the stochastic formation of differentially methylated regions in normal and cancerous tissues. Nat Genet. 2012 Nov;44(11):1207–14. doi:10.1038/ng.2442 PubMed PMID: 23064413.

28. Ntziachristos P, Tsirigos A, Van Vlierberghe P, Nedjic J, Trimarchi T, Flaherty MS, et al. Genetic inactivation of the polycomb repressive complex 2 in T cell acute lymphoblastic leukemia. Nat Med. 2012 Feb 6;18(2):298–301. doi:10.1038/nm.2651 PubMed PMID: 22237151; PubMed Central PMCID: PMC3274628.

29. Heery R, Schaefer MH. DNA methylation variation along the cancer epigenome and the identification of novel epigenetic driver events. Nucleic Acids Res. 2021 Dec 16;49(22):12692–705. doi:10.1093/nar/gkab1167

30. Parreno V, Martinez AM, Cavalli G. Mechanisms of Polycomb group protein function in cancer. Cell Res. 2022 Mar;32(3):231–53. doi:10.1038/s41422-021-00606-6 PubMed PMID: 35046519; PubMed Central PMCID: PMC8888700.

31. Chammas P, Mocavini I, Di Croce L. Engaging chromatin: PRC2 structure meets function. Br J Cancer. 2020 Feb;122(3):315–28. doi:10.1038/s41416-019-0615-2 PubMed PMID: 31708574; PubMed Central PMCID: PMC7000746.

32. Yücel AD, Gladyshev VN. Systemic epigenetic dysregulation as a driver of ageing and a therapeutic target. Nat Rev Mol Cell Biol. 2026 Jul;27(7):528–42. doi:10.1038/s41580-026-00958-0

33. Masika H, Ruppo S, Clark SJ, Bonder MJ, Von Meyenn F, Hecht M, et al. Cell-to-cell variability and gain of methylation at polycomb CpG islands as a hallmark of aging. Nat Commun. 2026 Jun 9;17(1):7318. doi:10.1038/s41467-026-74118-5

34. Camacho O, Koldobskiy MA, Reddy P, Oak A, Sun Y, Sherman K, et al. Convergence of aging- and rejuvenation-related epigenetic alterations on PRC2 targets. Mol Syst Biol. 2026 Feb 10;22(5):787–810. doi:10.1038/s44320-026-00195-9

35. Moqri M, Cipriano A, Simpson DJ, Rasouli S, Murty T, De Jong TA, et al. PRC2-AgeIndex as a universal biomarker of aging and rejuvenation. Nat Commun. 2024 Jul 16;15(1):5956. doi:10.1038/s41467-024-50098-2

36. Gao Y, Jones A, Fasching PA, Ruebner M, Beckmann MW, Widschwendter M, et al. The integrative epigenomic-transcriptomic landscape of ER positive breast cancer. Clin Epigenetics. 2015;7:126. doi:10.1186/s13148-015-0159-0 PubMed PMID: 26664652; PubMed Central PMCID: PMC4673726.

37. Tang W, Zhang F, Byun JS, Dorsey TH, Yfantis HG, Ajao A, et al. Population-specific Mutation Patterns in Breast Tumors from African American, European American, and Kenyan Patients. Cancer Res Commun. 2023 Nov 7;3(11):2244–55. doi:10.1158/2767-9764.CRC-23-0165 PubMed PMID: 37902422; PubMed Central PMCID: PMC10629394.

38. Dennis SR, Tsukioki T, Cottone G, Zhou W, Ganz PA, Sehl ME, et al. DNA methylation patterns in breast cancer, paired benign tissue from ipsilateral and contralateral breast, and healthy controls. Breast Cancer Res BCR. 2025 Jun 11;27(1):103. doi:10.1186/s13058-025-02057-y PubMed PMID: 40500734; PubMed Central PMCID: PMC12160127.

39. Muse ME, Titus AJ, Salas LA, Wilkins OM, Mullen C, Gregory KJ, et al. Enrichment of CpG island shore region hypermethylation in epigenetic breast field cancerization. Epigenetics. 2020 Oct;15(10):1093–106. doi:10.1080/15592294.2020.1747748 PubMed PMID: 32255732; PubMed Central PMCID: PMC7518670.

40. Johnson KC, Koestler DC, Fleischer T, Chen P, Jenson EG, Marotti JD, et al. DNA methylation in ductal carcinoma in situ related with future development of invasive breast cancer. Clin Epigenetics. 2015;7(1):75. doi:10.1186/s13148-015-0094-0 PubMed PMID: 26213588; PubMed Central PMCID: PMC4514996.

41. Schultz BM, Ayeotan O, Rana JN, Litwin S, Basickes S, Sapienza C, et al. Abnormal epigenetic aging of epigenetic outliers in normal colon mucosa from colorectal cancer patients. Sci Rep. 2026 Jan 7;16(1):4022. doi:10.1038/s41598-025-34035-x PubMed PMID: 41501347; PubMed Central PMCID: PMC12855823.

42. Luo Y, Wong CJ, Kaz AM, Dzieciatkowski S, Carter KT, Morris SM, et al. Differences in DNA methylation signatures reveal multiple pathways of progression from adenoma to colorectal cancer. Gastroenterology. 2014 Aug;147(2):418–429.e8. doi:10.1053/j.gastro.2014.04.039 PubMed PMID: 24793120; PubMed Central PMCID: PMC4107146.

43. Murgas KA, Ma Y, Shahidi LK, Mukherjee S, Allen AS, Shibata D, et al. A Bayesian hierarchical model to estimate DNA methylation conservation in colorectal tumors. Bioinformatics. 2021 Dec 22;38(1):22–9. doi:10.1093/bioinformatics/btab637 PubMed PMID: 34487148; PubMed Central PMCID: PMC10060692.

44. Karlsson A, Jönsson M, Lauss M, Brunnström H, Jönsson P, Borg Å, et al. Genome-wide DNA methylation analysis of lung carcinoma reveals one neuroendocrine and four adenocarcinoma epitypes associated with patient outcome. Clin Cancer Res Off J Am Assoc Cancer Res. 2014 Dec 1;20(23):6127–40. doi:10.1158/1078-0432.CCR-14-1087 PubMed PMID: 25278450.

45. Teixeira VH, Pipinikas CP, Pennycuick A, Lee-Six H, Chandrasekharan D, Beane J, et al. Deciphering the genomic, epigenomic, and transcriptomic landscapes of pre-invasive lung cancer lesions. Nat Med. 2019 Mar;25(3):517–25. doi:10.1038/s41591-018-0323-0 PubMed PMID: 30664780; PubMed Central PMCID: PMC7614970.

46. Kuramoto J, Arai E, Tian Y, Funahashi N, Hiramoto M, Nammo T, et al. Genome-wide DNA methylation analysis during non-alcoholic steatohepatitis-related multistage hepatocarcinogenesis: comparison with hepatitis virus-related carcinogenesis. Carcinogenesis. 2017 Mar 1;38(3):261–70. doi:10.1093/carcin/bgx005 PubMed PMID: 28426876; PubMed Central PMCID: PMC5862314.

47. Aref-Eshghi E, Schenkel LC, Ainsworth P, Lin H, Rodenhiser DI, Cutz JC, et al. Genomic DNA Methylation-Derived Algorithm Enables Accurate Detection of Malignant Prostate Tissues. Front Oncol. 2018;8:100. doi:10.3389/fonc.2018.00100 PubMed PMID: 29740534; PubMed Central PMCID: PMC5925605.

48. Silva R, Moran B, Baird AM, O’Rourke CJ, Finn SP, McDermott R, et al. Longitudinal analysis of individual cfDNA methylome patterns in metastatic prostate cancer. Clin Epigenetics. 2021 Aug 28;13(1):168. doi:10.1186/s13148-021-01155-w PubMed PMID: 34454584; PubMed Central PMCID: PMC8403420.

49. Luebeck EG, Curtius K, Hazelton WD, Maden S, Yu M, Thota PN, et al. Identification of a key role of widespread epigenetic drift in Barrett’s esophagus and esophageal adenocarcinoma. Clin Epigenetics. 2017;9:113. doi:10.1186/s13148-017-0409-4 PubMed PMID: 29046735; PubMed Central PMCID: PMC5644061.

50. Krause L, Nones K, Loffler KA, Nancarrow D, Oey H, Tang YH, et al. Identification of the CIMP-like subtype and aberrant methylation of members of the chromosomal segregation and spindle assembly pathways in esophageal adenocarcinoma. Carcinogenesis. 2016 Apr;37(4):356–65. doi:10.1093/carcin/bgw018 PubMed PMID: 26905591; PubMed Central PMCID: PMC4806711.

51. Huang KK, Ramnarayanan K, Zhu F, Srivastava S, Xu C, Tan ALK, et al. Genomic and Epigenomic Profiling of High-Risk Intestinal Metaplasia Reveals Molecular Determinants of Progression to Gastric Cancer. Cancer Cell. 2018 Jan 8;33(1):137–150.e5. doi:10.1016/j.ccell.2017.11.018 PubMed PMID: 29290541.

52. Goeppert B, Stichel D, Toth R, Fritzsche S, Loeffler MA, Schlitter AM, et al. Integrative analysis reveals early and distinct genetic and epigenetic changes in intraductal papillary and tubulopapillary cholangiocarcinogenesis. Gut. 2022 Feb;71(2):391–401. doi:10.1136/gutjnl-2020-322983 PubMed PMID: 33468537; PubMed Central PMCID: PMC8762040.

53. Németh CG, Röcken C, Siebert R, Wiltfang J, Ammerpohl O, Gassling V. Recurrent chromosomal and epigenetic alterations in oral squamous cell carcinoma and its putative premalignant condition oral lichen planus. PloS One. 2019;14(4):e0215055. doi:10.1371/journal.pone.0215055 PubMed PMID: 30964915; PubMed Central PMCID: PMC6456184.

54. Bian S, Hou Y, Zhou X, Li X, Yong J, Wang Y, et al. Single-cell multiomics sequencing and analyses of human colorectal cancer. Science. 2018 Nov 30;362(6418):1060–3. doi:10.1126/science.aao3791 PubMed PMID: 30498128.

55. Kelly K, Scherer M, Braun MM, Lutsik P, Plass C. EpiCHAOS: a metric to quantify epigenomic heterogeneity in single-cell data. Genome Biol. 2024 Dec 3;25(1):305. doi:10.1186/s13059-024-03446-w PubMed PMID: 39623476; PubMed Central PMCID: PMC11613708.

56. Sheffield NC, Bock C. LOLA: enrichment analysis for genomic region sets and regulatory elements in R and Bioconductor. Bioinformatics. 2016 Feb 15;32(4):587–9. doi:10.1093/bioinformatics/btv612

57. The ENCODE Project Consortium. An integrated encyclopedia of DNA elements in the human genome. Nature. 2012 Sep;489(7414):57–74. doi:10.1038/nature11247

58. Roadmap Epigenomics Consortium, Kundaje A, Meuleman W, Ernst J, Bilenky M, Yen A, et al. Integrative analysis of 111 reference human epigenomes. Nature. 2015 Feb 19;518(7539):317–30. doi:10.1038/nature14248

59. Liu T, Ortiz JA, Taing L, Meyer CA, Lee B, Zhang Y, et al. Cistrome: an integrative platform for transcriptional regulation studies. Genome Biol. 2011 Aug 22;12(8):R83. doi:10.1186/gb-2011-12-8-r83

60. Kagda MS, Lam B, Litton C, Small C, Sloan CA, Spragins E, et al. Data navigation on the ENCODE portal. Nat Commun. 2025 Oct 30;16(1):9592. doi:10.1038/s41467-025-64343-9 PubMed PMID: 41168159; PubMed Central PMCID: PMC12575607.

61. Gao C, Amador C, Walker RM, Campbell A, Madden RA, Adams MJ, et al. Phenome-wide analyses identify an association between the parent-of-origin effects dependent methylome and the rate of aging in humans. Genome Biol. 2023 May 15;24(1):117. doi:10.1186/s13059-023-02953-6

62. Ernst J, Kellis M. ChromHMM: automating chromatin-state discovery and characterization. Nat Methods. 2012 Mar;9(3):215–6. doi:10.1038/nmeth.1906

63. Dale RK, Pedersen BS, Quinlan AR. Pybedtools: a flexible Python library for manipulating genomic datasets and annotations. Bioinformatics. 2011 Dec 15;27(24):3423–4. doi:10.1093/bioinformatics/btr539

64. Corces MR, Granja JM, Shams S, Louie BH, Seoane JA, Zhou W, et al. The chromatin accessibility landscape of primary human cancers. Science. 2018 Oct 26;362(6413):eaav1898. doi:10.1126/science.aav1898 PubMed PMID: 30361341; PubMed Central PMCID: PMC6408149.

65. Dobin A, Davis CA, Schlesinger F, Drenkow J, Zaleski C, Jha S, et al. STAR: ultrafast universal RNA-seq aligner. Bioinformatics. 2013 Jan 1;29(1):15–21. doi:10.1093/bioinformatics/bts635

66. Lawrence M, Gentleman R, Carey V. rtracklayer: an R package for interfacing with genome browsers. Bioinformatics. 2009 Jul 15;25(14):1841–2. doi:10.1093/bioinformatics/btp328 PubMed PMID: 19468054; PubMed Central PMCID: PMC2705236.

67. Lawrence M, Huber W, Pagès H, Aboyoun P, Carlson M, Gentleman R, et al. Software for Computing and Annotating Genomic Ranges. Prlic A, editor. PLoS Comput Biol. 2013 Aug 8;9(8):e1003118. doi:10.1371/journal.pcbi.1003118

68. Marc Carlson BPMO [Cre. TxDb.Hsapiens.UCSC.hg19.knownGene [Internet]. Bioconductor; 2017 [cited 2026 Oct 5]. Available from: https://bioconductor.org/packages/TxDb.Hsapiens.UCSC.hg19.knownGene doi:10.18129/B9.BIOC.TXDB.HSAPIENS.UCSC.HG19.KNOWNGENE

69. Hervé Pagès MC. AnnotationDbi [Internet]. Bioconductor; 2017 [cited 2026 Oct 5]. Available from: https://bioconductor.org/packages/AnnotationDbi doi:10.18129/B9.BIOC.ANNOTATIONDBI

70. Davidson-Pilon C. lifelines: survival analysis in Python. J Open Source Softw. 2019 Aug 4;4(40):1317. doi:10.21105/joss.01317

