## Supplementary Figures for "PRC2-binding regions are hotspots of DNA methylation entropy in pre-cancer and cancer"

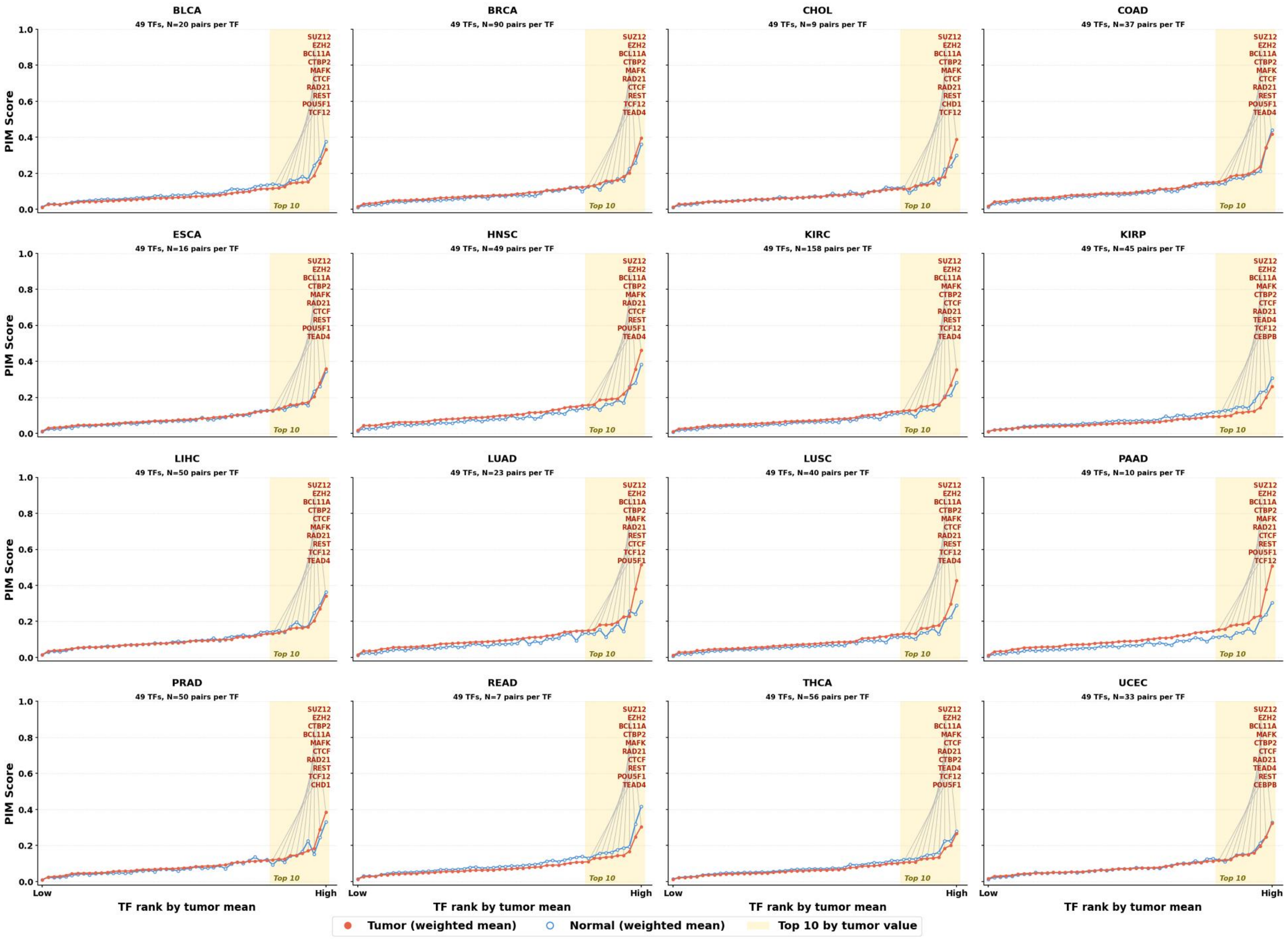
BLCA
49 TFs, N=20 pairs per TF
SUZ12 EZH2 BCL11A CTBP2 MAFK CTCF RAD21 REST POU5F1 TCF12
BRCA
49 TFs, N=90 pairs per TF
SUZ12 EZH2 BCL11A CTBP2 MAFK RAD21 CTCF REST TCF12 TEAD4
CHOL
49 TFs, N=9 pairs per TF
SUZ12 EZH2 BCL11A CTBP2 MAFK CTCF RAD21 REST CHD1 TCF12
COAD
49 TFs, N=37 pairs per TF
SUZ12 EZH2 BCL11A CTBP2 MAFK CTCF RAD21 REST POU5F1 TEAD4
ESCA
49 TFs, N=16 pairs per TF
SUZ12 EZH2 BCL11A CTBP2 MAFK RAD21 CTCF REST POU5F1 TEAD4
HNSC
49 TFs, N=49 pairs per TF
SUZ12 EZH2 BCL11A CTBP2 MAFK RAD21 CTCF REST POU5F1 TEAD4
KIRC
49 TFs, N=158 pairs per TF
SUZ12 EZH2 BCL11A MAFK CTBP2 CTCF RAD21 REST TCF12 TEAD4
KIRP
49 TFs, N=45 pairs per TF
SUZ12 EZH2 BCL11A MAFK CTBP2 CTCF RAD21 TEAD4 TCF12 CEBPB
LIHC
49 TFs, N=50 pairs per TF
SUZ12 EZH2 BCL11A CTBP2 CTCF MAFK RAD21 REST TCF12 TEAD4
LUAD
49 TFs, N=23 pairs per TF
SUZ12 EZH2 BCL11A CTBP2 MAFK RAD21 REST CTCF TCF12 POU5F1
LUSC
49 TFs, N=40 pairs per TF
SUZ12 EZH2 BCL11A CTBP2 MAFK CTCF RAD21 REST TCF12 TEAD4
PAAD
49 TFs, N=10 pairs per TF
SUZ12 EZH2 BCL11A CTBP2 MAFK RAD21 CTCF REST POU5F1 TCF12
PRAD
49 TFs, N=50 pairs per TF
SUZ12 EZH2 CTBP2 BCL11A MAFK CTCF RAD21 REST TCF12 CHD1
READ
49 TFs, N=7 pairs per TF
SUZ12 EZH2 BCL11A CTBP2 MAFK RAD21 CTCF REST POU5F1 TEAD4
THCA
49 TFs, N=56 pairs per TF
SUZ12 EZH2 BCL11A MAFK CTCF RAD21 CTBP2 TEAD4 TCF12 POU5F1
UCEC
49 TFs, N=33 pairs per TF
SUZ12 EZH2 BCL11A MAFK CTBP2 CTCF RAD21 TEAD4 REST CEBPB
Top 10
PIM Score
0.0 0.2 0.4 0.6 0.8 1.0
Low High
TF rank by tumor mean
Tumor (weighted mean)
Normal (weighted mean)
Top 10 by tumor value

1

**Supplementary Figure 1. Cancer type–specific PIM scores across transcription factor binding regions.** PIM scores were quantified within binding regions of 49 transcription factors (TFs) across 16 cancer types using matched tumor–normal samples. TFs are ranked from low to high according to the tumor weighted mean PIM score within each cancer type. Tumor values are shown as filled red circles and matched normal values as open blue circles. The shaded area denotes the 10 TFs with the highest tumor-associated PIM scores, which are labeled within each panel. The number of matched tumor–normal pairs analyzed per TF is indicated above each cancer-specific panel.

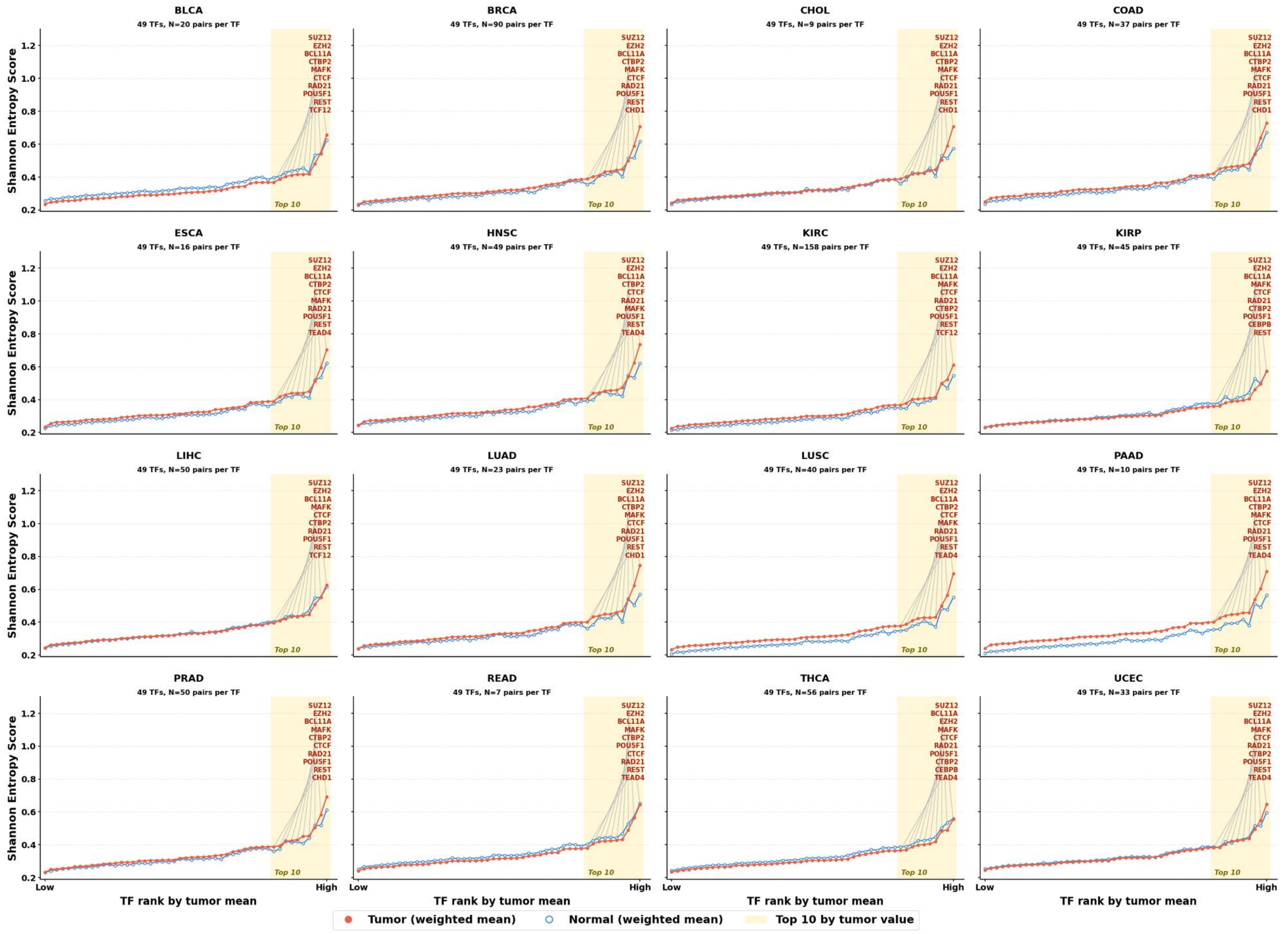
BLCA
49 TFs, N=20 pairs per TF
SUZ12 EZH2 BCL11A CTBP2 MAFK CTCF RAD21 POU5F1 REST TCF12
BRCA
49 TFs, N=90 pairs per TF
SUZ12 EZH2 BCL11A CTBP2 MAFK CTCF RAD21 POU5F1 REST CHD1
CHOL
49 TFs, N=9 pairs per TF
SUZ12 EZH2 BCL11A CTBP2 MAFK CTCF RAD21 POU5F1 REST CHD1
COAD
49 TFs, N=37 pairs per TF
SUZ12 EZH2 BCL11A CTBP2 MAFK CTCF RAD21 POU5F1 REST CHD1
ESCA
49 TFs, N=16 pairs per TF
SUZ12 EZH2 BCL11A CTBP2 CTCF MAFK RAD21 POU5F1 REST TEAD4
HNSC
49 TFs, N=49 pairs per TF
SUZ12 EZH2 BCL11A CTBP2 CTCF RAD21 MAFK POU5F1 REST TEAD4
KIRC
49 TFs, N=158 pairs per TF
SUZ12 EZH2 BCL11A MAFK CTCF RAD21 CTBP2 POU5F1 REST TCF12
KIRP
49 TFs, N=45 pairs per TF
SUZ12 EZH2 BCL11A MAFK CTCF RAD21 CTBP2 POU5F1 CEBPB REST
LIHC
49 TFs, N=50 pairs per TF
SUZ12 EZH2 BCL11A MAFK CTCF CTBP2 RAD21 POU5F1 REST TCF12
LUAD
49 TFs, N=23 pairs per TF
SUZ12 EZH2 BCL11A CTBP2 MAFK CTCF RAD21 POU5F1 REST CHD1
LUSC
49 TFs, N=40 pairs per TF
SUZ12 EZH2 BCL11A CTBP2 CTCF MAFK RAD21 POU5F1 REST TEAD4
PAAD
49 TFs, N=10 pairs per TF
SUZ12 EZH2 BCL11A CTBP2 MAFK CTCF RAD21 POU5F1 REST TEAD4
PRAD
49 TFs, N=50 pairs per TF
SUZ12 EZH2 BCL11A MAFK CTBP2 CTCF RAD21 POU5F1 REST CHD1
READ
49 TFs, N=7 pairs per TF
SUZ12 EZH2 BCL11A MAFK CTBP2 POU5F1 CTCF RAD21 REST TEAD4
THCA
49 TFs, N=56 pairs per TF
SUZ12 BCL11A EZH2 MAFK CTCF RAD21 POU5F1 CTBP2 CEBPB TEAD4
UCEC
49 TFs, N=33 pairs per TF
SUZ12 EZH2 BCL11A MAFK CTCF RAD21 CTBP2 POU5F1 REST TEAD4
Top 10
Shannon Entropy Score
0.2 0.4 0.6 0.8 1.0 1.2
Low High
TF rank by tumor mean
Tumor (weighted mean)
Normal (weighted mean)
Top 10 by tumor value

**Supplementary Figure 2. Cancer type–specific Shannon entropy scores across transcription factor binding regions.** Shannon entropy scores were quantified within binding regions of 49 transcription factors (TFs) across 16 cancer types using matched tumor–normal samples. TFs are ranked from low to high according to the tumor weighted mean Shannon entropy score within each cancer type. Tumor values are shown as filled red circles and matched normal values as open blue circles. The shaded area denotes the 10 TFs with the highest tumor-associated Shannon entropy scores, which are labeled within each panel. The number of matched tumor–normal pairs analyzed per TF is indicated above each cancer-specific panel.

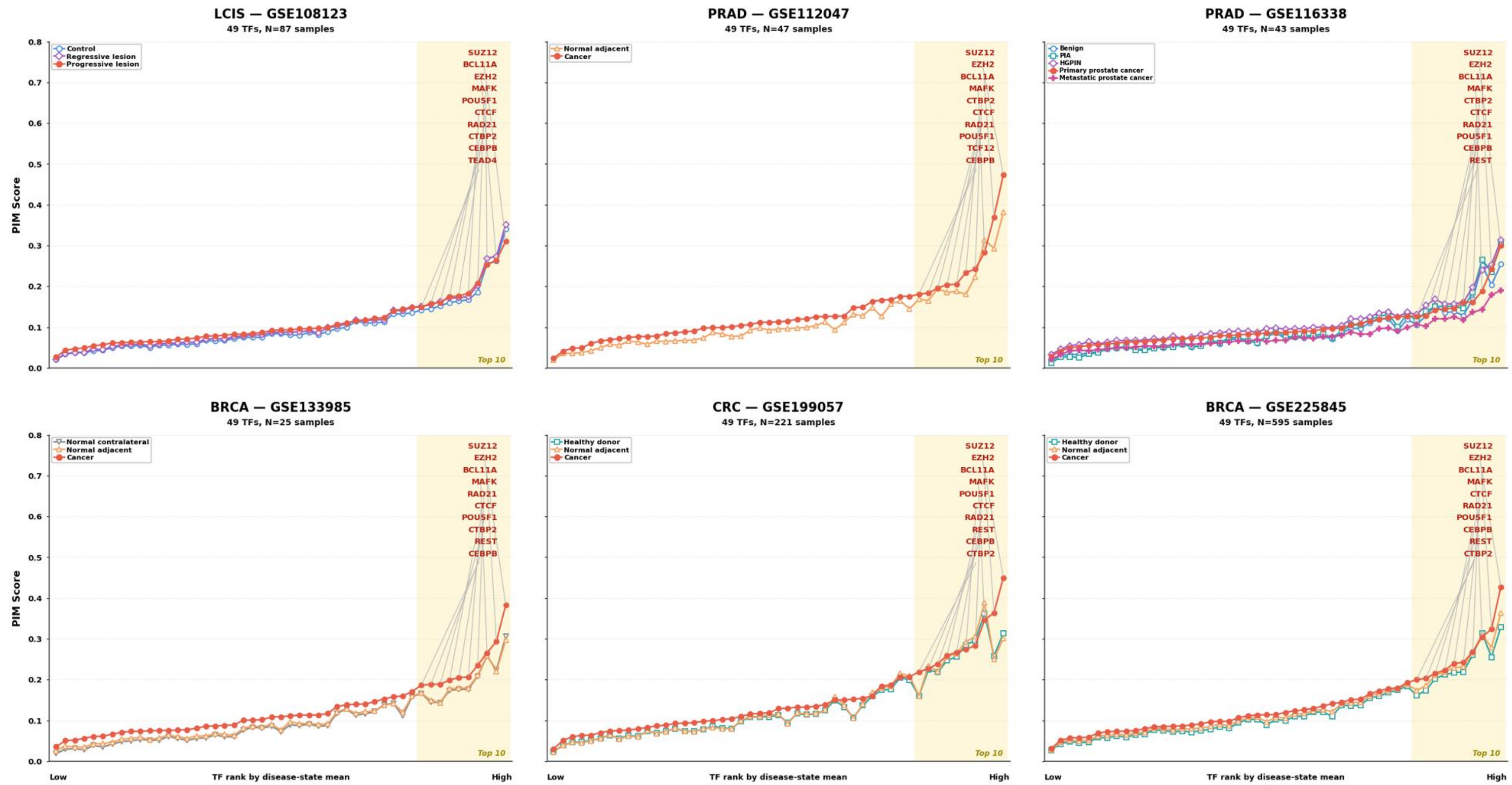


**Supplementary Figure 3. PIM scores across transcription factor binding regions in independent validation cohorts spanning normal, premalignant, progressive, regressive, and malignant tissue states.** PIM scores were quantified within binding regions of 49 transcription factors (TFs) in six independent GEO datasets: pre-invasive lung squamous lesions (LCIS; GSE108123), prostate cancer (PRAD; GSE112047 and GSE116338), breast cancer (BRCA; GSE133985 and GSE225845), and colorectal cancer (CRC; GSE199057). TFs are ranked from low to high according to the mean PIM score of the disease-state group within each dataset. The shaded area denotes the 10 TFs with the highest PIM scores, which are labeled within each panel. Tissue and disease-state groups, together with total sample numbers, are indicated within the corresponding panels.

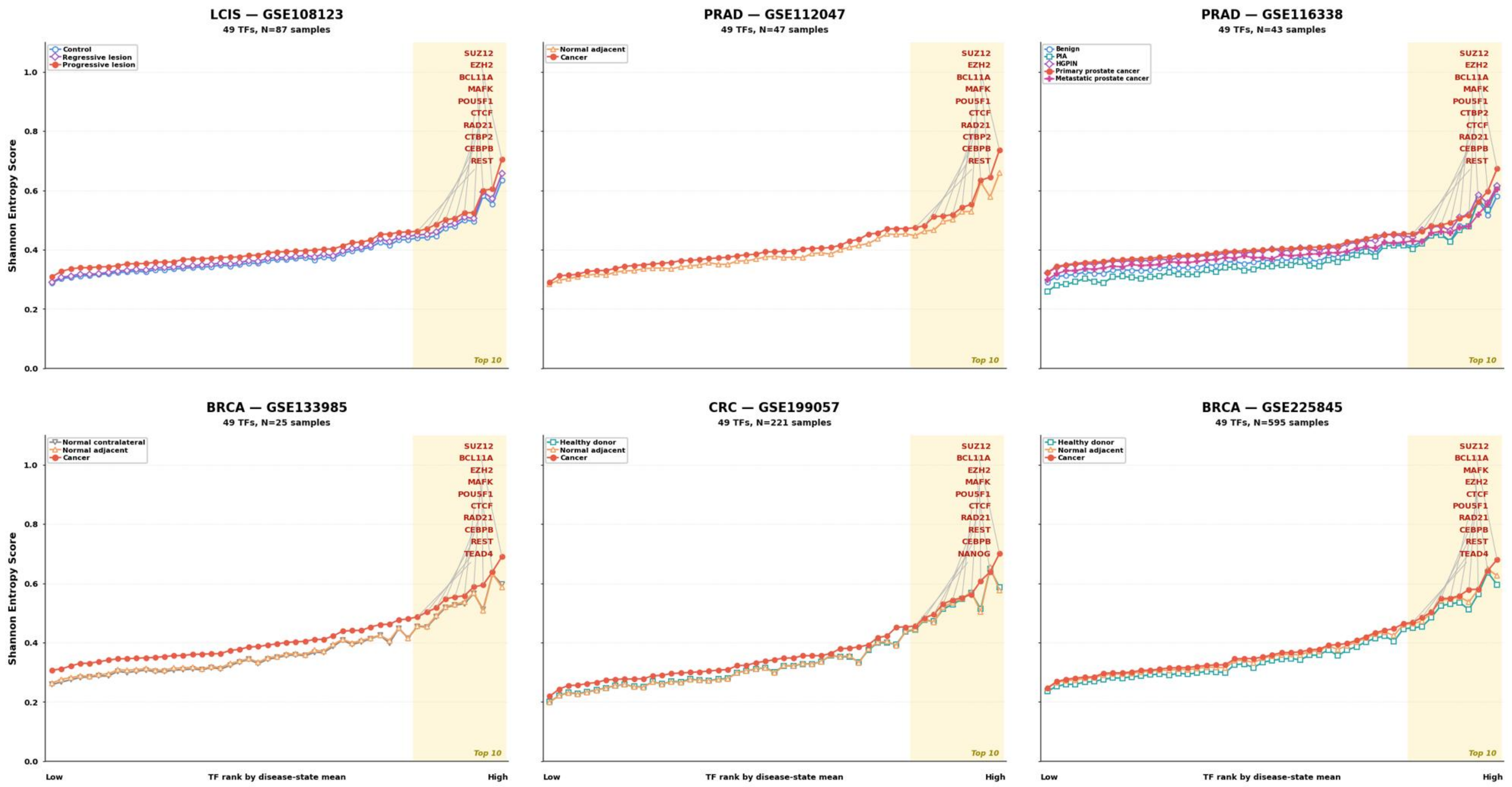


**Supplementary Figure 4. Shannon entropy scores across transcription factor binding regions in independent validation cohorts spanning normal, premalignant, progressive, regressive, and malignant tissue states.** Shannon entropy scores were quantified within binding regions of 49 transcription factors (TFs) in six independent GEO datasets: pre-invasive lung squamous lesions (LCIS; GSE108123), prostate cancer (PRAD; GSE112047 and GSE116338), breast cancer (BRCA; GSE133985 and GSE225845), and colorectal cancer (CRC; GSE199057). TFs are ranked from low to high according to the mean Shannon entropy score of the disease-state group within each dataset. The shaded area denotes the 10 TFs with the highest Shannon entropy scores, which are labeled within each panel. Tissue and disease-state groups, together with total sample numbers, are indicated within the corresponding panels.

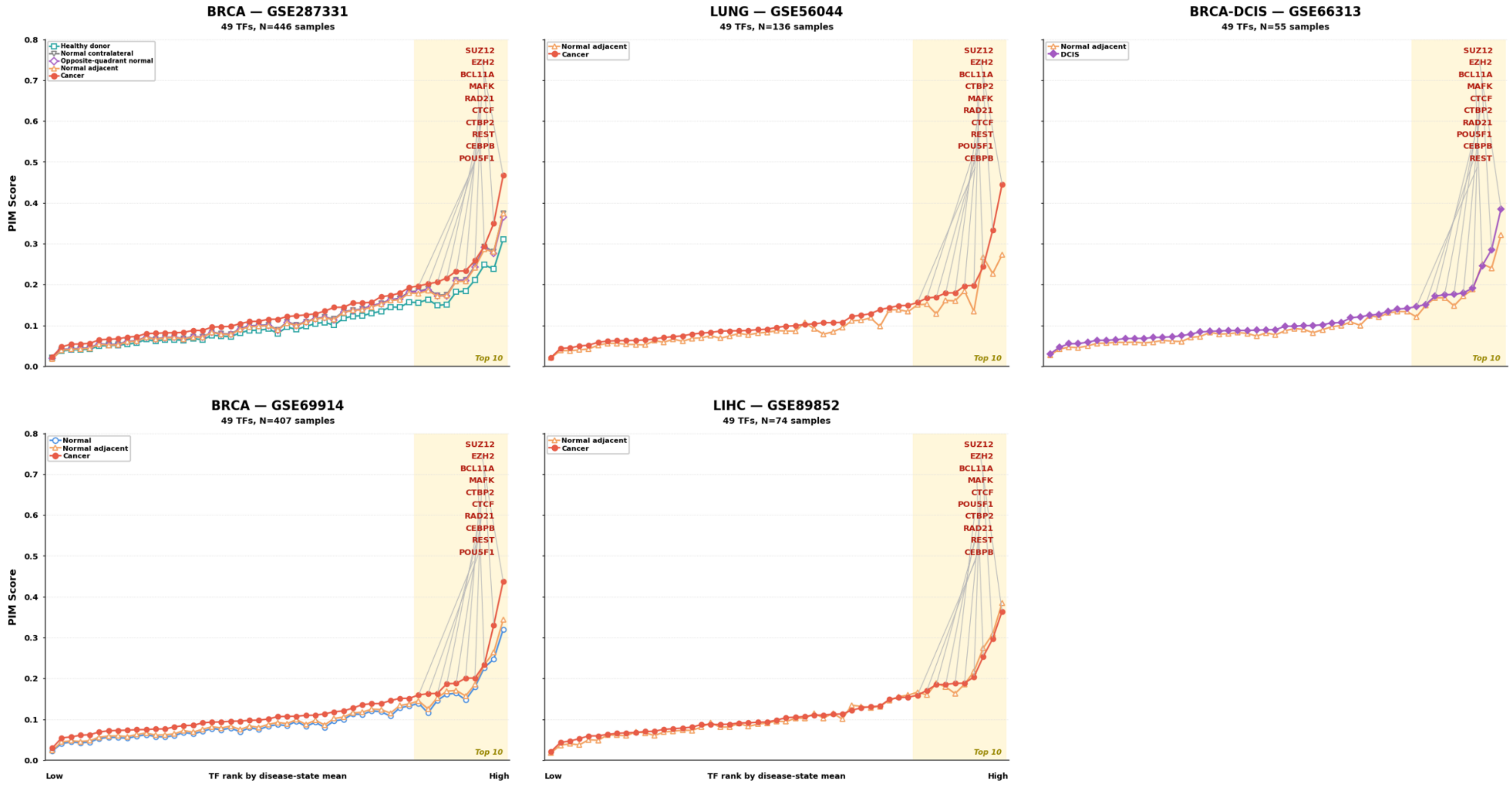


**Supplementary Figure 5. PIM scores across transcription factor binding regions in additional independent validation cohorts.** PIM scores were quantified within binding regions of 49 transcription factors (TFs) in five independent GEO datasets: breast cancer and breast field-effect cohorts (BRCA; GSE287331 and GSE69914), lung cancer (GSE56044), breast ductal carcinoma in situ (BRCA-DCIS; GSE66313), and hepatocellular carcinoma (LIHC; GSE89852). TFs are ranked from low to high according to the mean PIM score of the disease-state group within each dataset. The shaded area denotes the 10 TFs with the highest PIM scores, which are labeled within each panel. Tissue and disease-state groups, together with total sample numbers, are indicated within the corresponding panels.

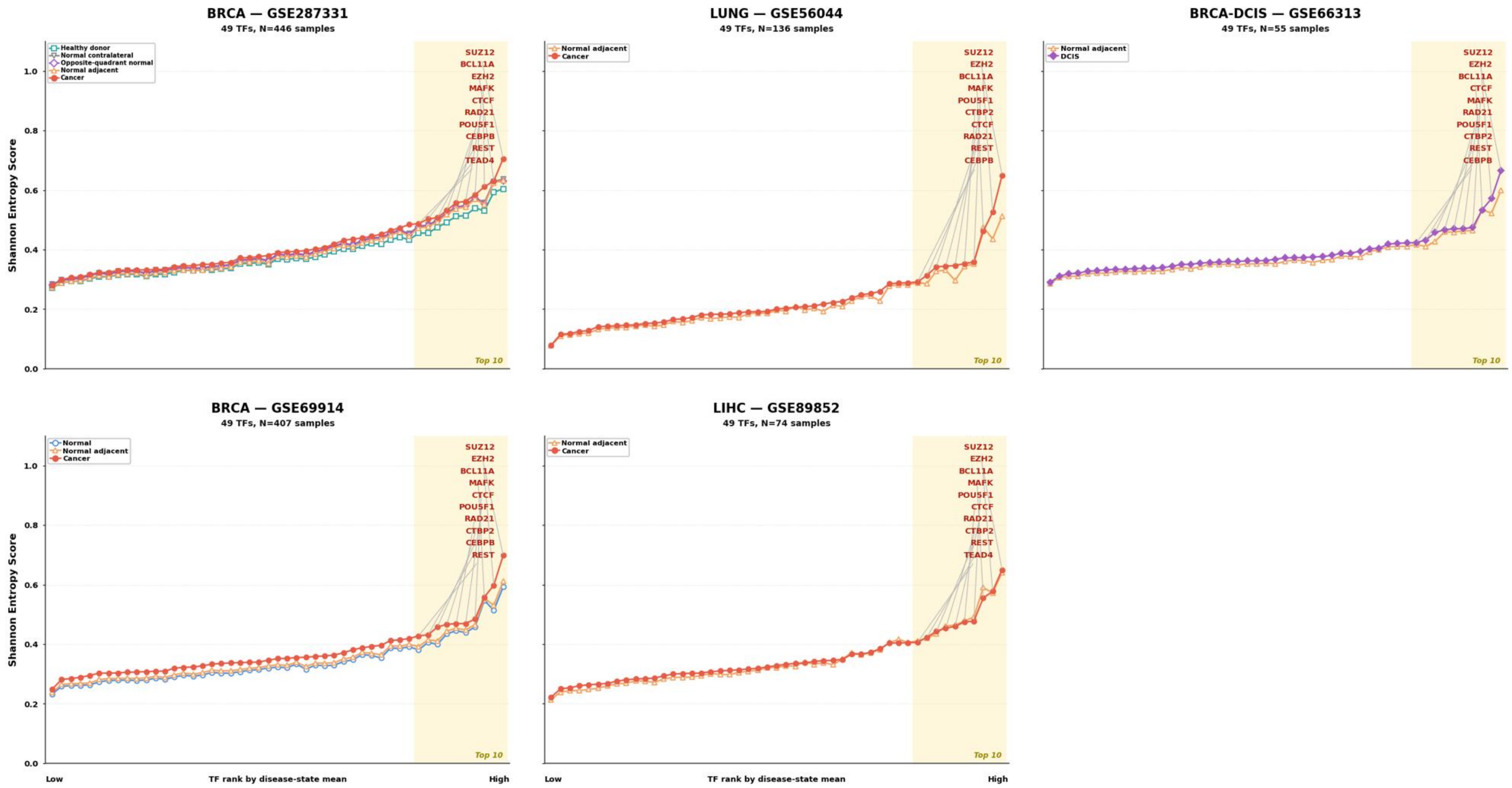


**Supplementary Figure 6. Shannon entropy scores across transcription factor binding regions in additional independent validation cohorts.** Shannon entropy scores were quantified within binding regions of 49 transcription factors (TFs) in five independent GEO datasets: breast cancer and breast field-effect cohorts (BRCA; GSE287331 and GSE69914), lung cancer (GSE56044), breast ductal carcinoma in situ (BRCA-DCIS; GSE66313), and hepatocellular carcinoma (LIHC; GSE89852). TFs are ranked from low to high according to the mean Shannon entropy score of the disease-state group within each dataset. The shaded area denotes the 10 TFs with the highest Shannon entropy scores, which are labeled within each panel. Tissue and disease-state groups, together with total sample numbers, are indicated within the corresponding panels.

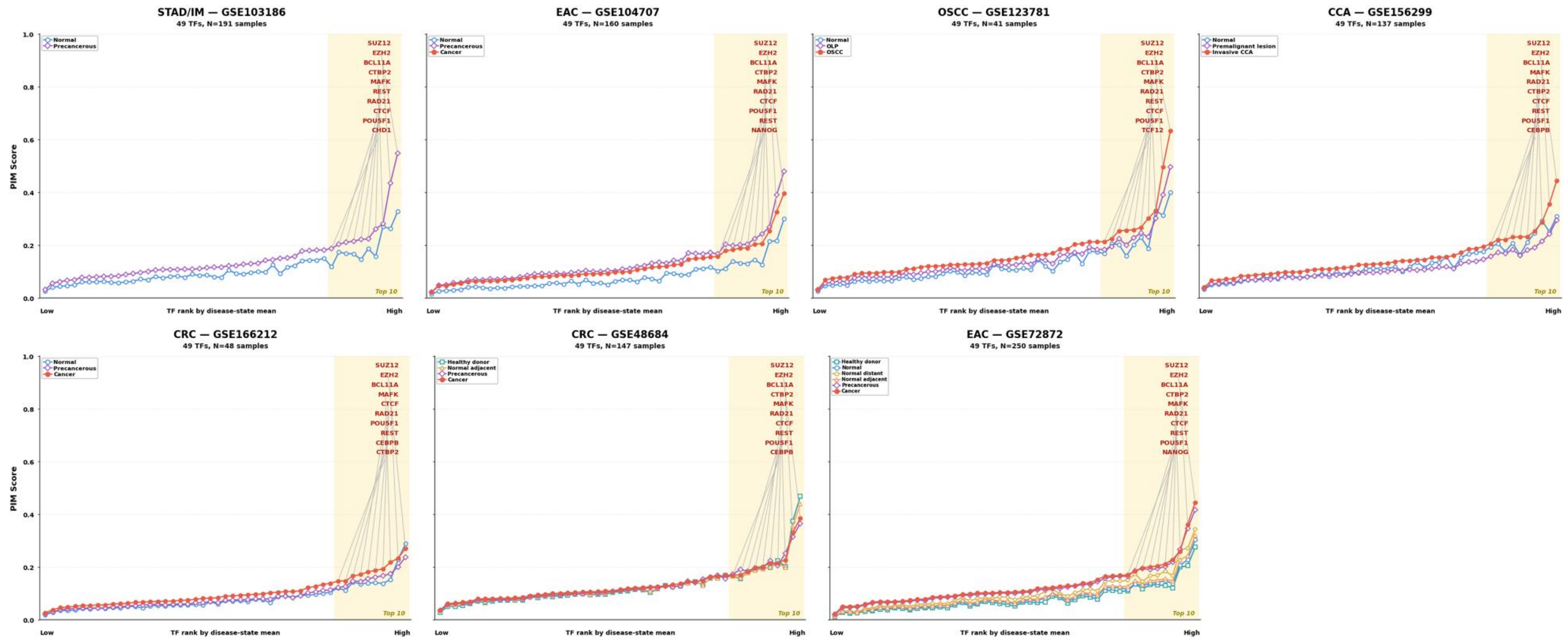


**Supplementary Figure 7. PIM scores across transcription factor binding regions in independent validation cohorts spanning premalignant and malignant tissue states.** PIM scores were quantified within binding regions of 49 transcription factors (TFs) in seven independent GEO datasets: gastric intestinal metaplasia and gastric cancer progression (STAD/IM; GSE103186), esophageal adenocarcinoma (EAC; GSE104707 and GSE72872), oral squamous cell carcinoma (OSCC; GSE123781), cholangiocarcinoma and premalignant lesions (CCA; GSE156299), and colorectal cancer and adenoma–carcinoma progression (CRC; GSE166212 and GSE48684). TFs are ranked from low to high according to the mean PIM score of the disease-state group within each dataset. The shaded area denotes the 10 TFs with the highest PIM scores, which are labeled within each panel. Tissue and disease-state groups, together with total sample numbers, are indicated within the corresponding panels.

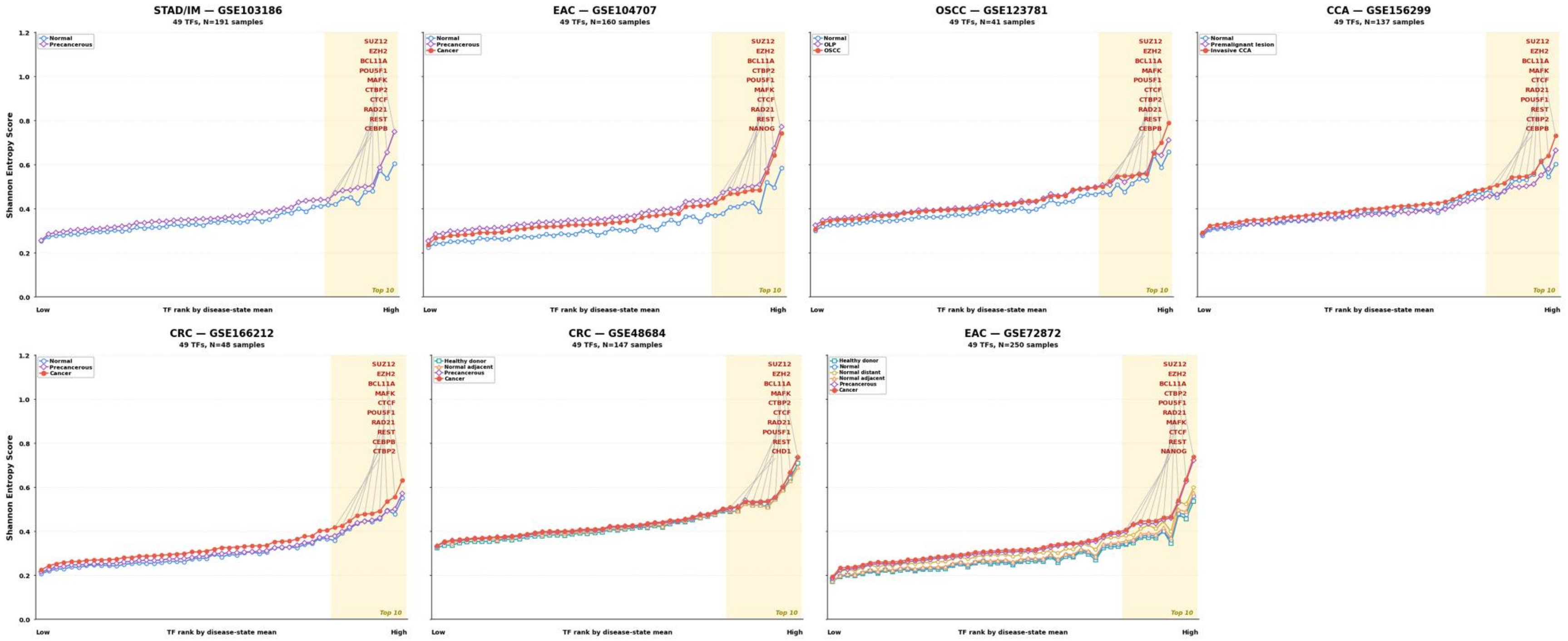


**Supplementary Figure 8. Shannon entropy scores across transcription factor binding regions in independent validation cohorts spanning premalignant and malignant tissue states.** Shannon entropy scores were quantified within binding regions of 49 transcription factors (TFs) in seven independent GEO datasets: gastric intestinal metaplasia and gastric cancer progression (STAD/IM; GSE103186), esophageal adenocarcinoma (EAC; GSE104707 and GSE72872), oral squamous cell carcinoma (OSCC; GSE123781), cholangiocarcinoma and premalignant lesions (CCA; GSE156299), and colorectal cancer and adenoma–carcinoma progression (CRC; GSE166212 and GSE48684). TFs are ranked from low to high according to the mean Shannon entropy score of the disease-state group within each dataset. The shaded area denotes the 10 TFs with the highest Shannon entropy scores, which are labeled within each panel. Tissue and disease-state groups, together with total sample numbers, are indicated within the corresponding panels.

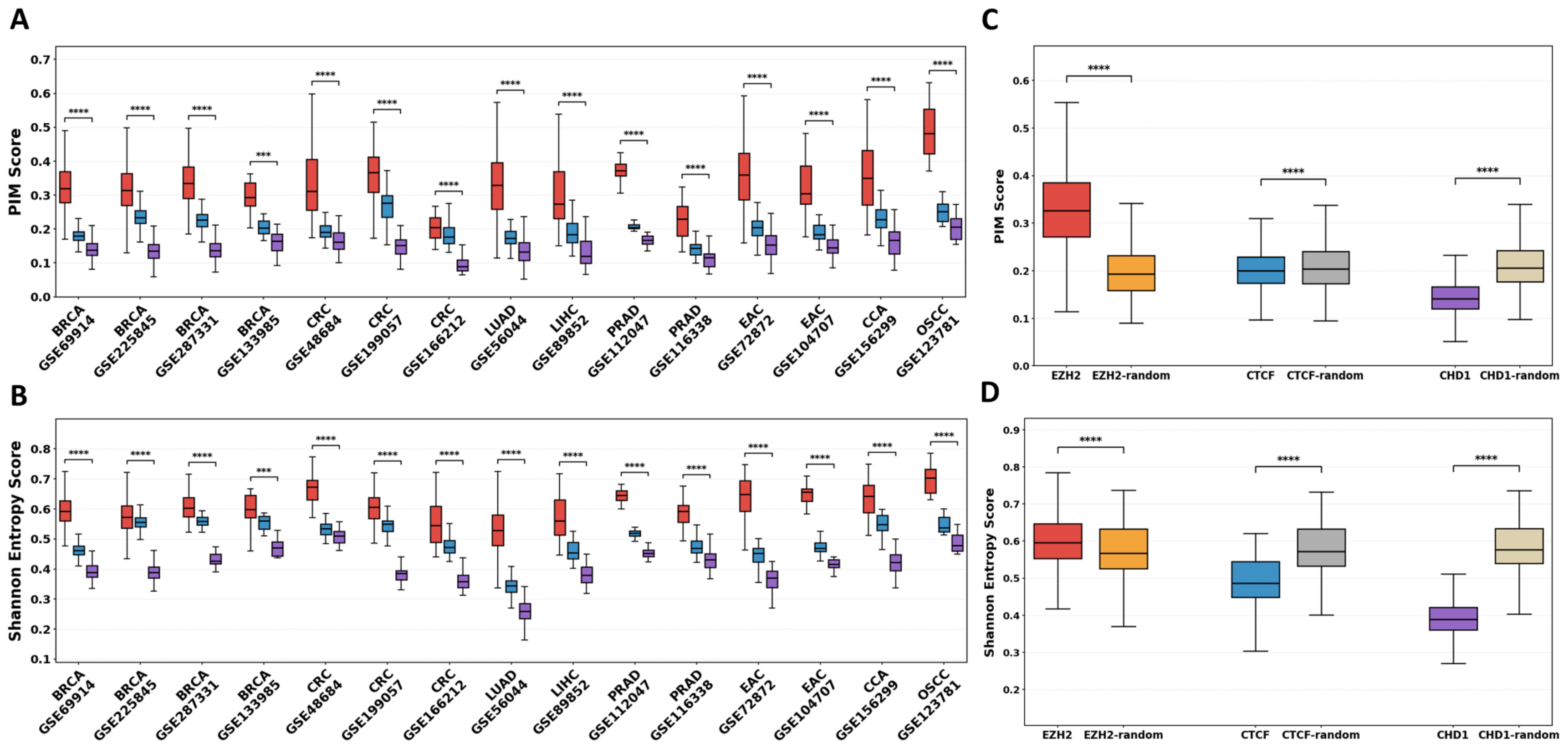


**Supplementary Figure 9. DNA methylation variability across EZH2-, CTCF-, and CHD1-binding regions in independent validation cohorts. A)** PIM scores within EZH2-, CTCF-, and CHD1-binding regions across the indicated independent validation datasets. **B)** Pooled comparison of PIM scores within EZH2-, CTCF-, and CHD1-binding regions and their corresponding matched random control regions. **C)** Shannon entropy scores within EZH2-, CTCF-, and CHD1-binding regions across the indicated validation datasets. **D)** Pooled comparison of Shannon entropy scores within EZH2-, CTCF-, and CHD1-binding regions and their corresponding matched random control regions. Boxplots show the median, interquartile range, and whiskers. Statistical significance is indicated above the corresponding comparisons (***P < 0.001; ****P < 0.0001).

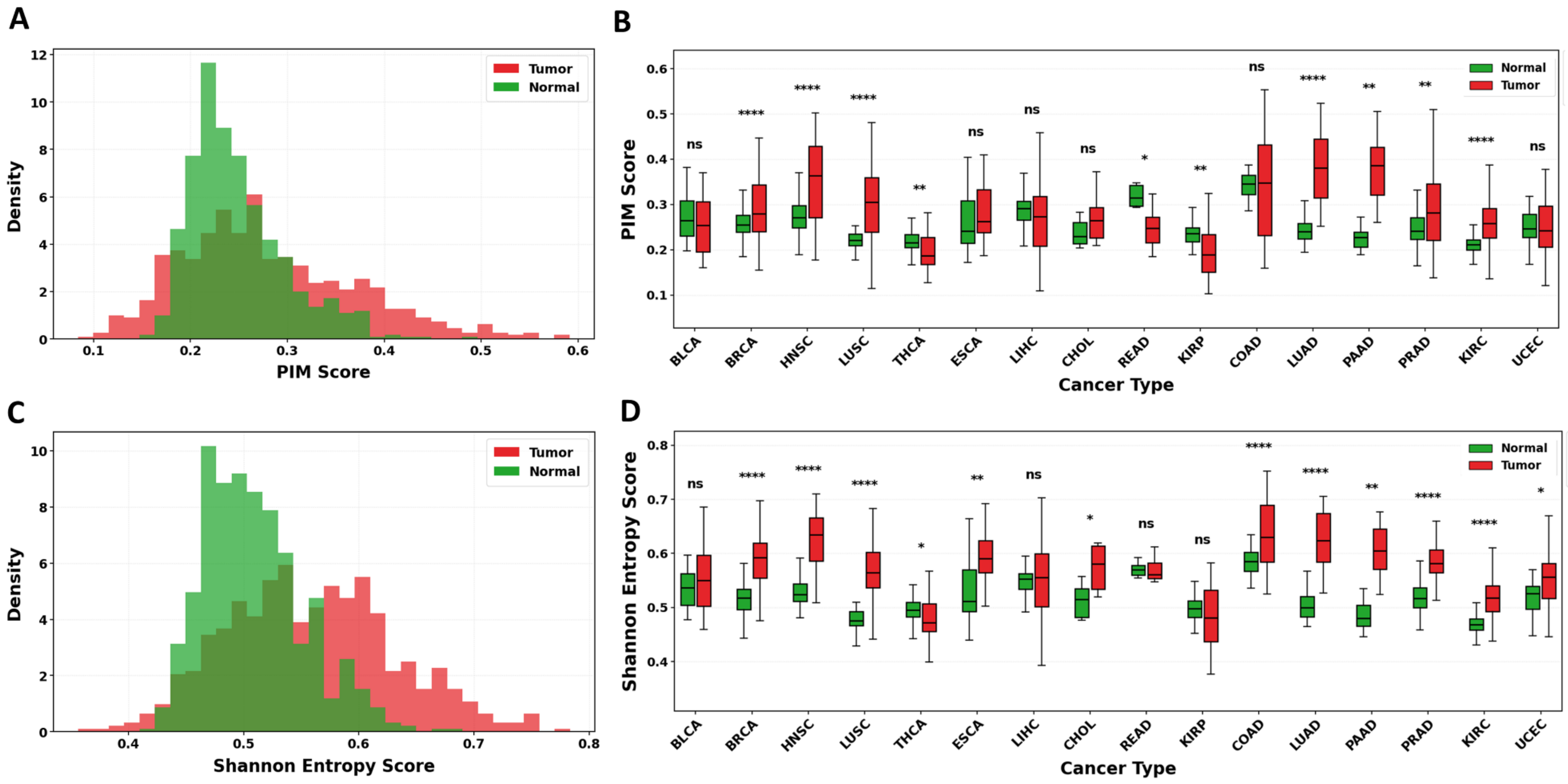


**Supplementary Figure 10. Distribution of PIM and Shannon entropy scores in matched normal and tumor tissues across cancer types. A)** Density distribution of PIM scores in pooled matched normal and tumor samples. **B)** PIM scores in normal and tumor samples across the indicated cancer types. **C)** Density distribution of Shannon entropy scores in pooled matched normal and tumor samples. **D)** Shannon entropy scores in normal and tumor samples across the indicated cancer types. Boxplots show the median, interquartile range, and whiskers. Statistical significance is indicated above the corresponding tumor–normal comparisons (ns, not significant; *P* < 0.05; **P** < 0.01; ** **P** < 0.0001).

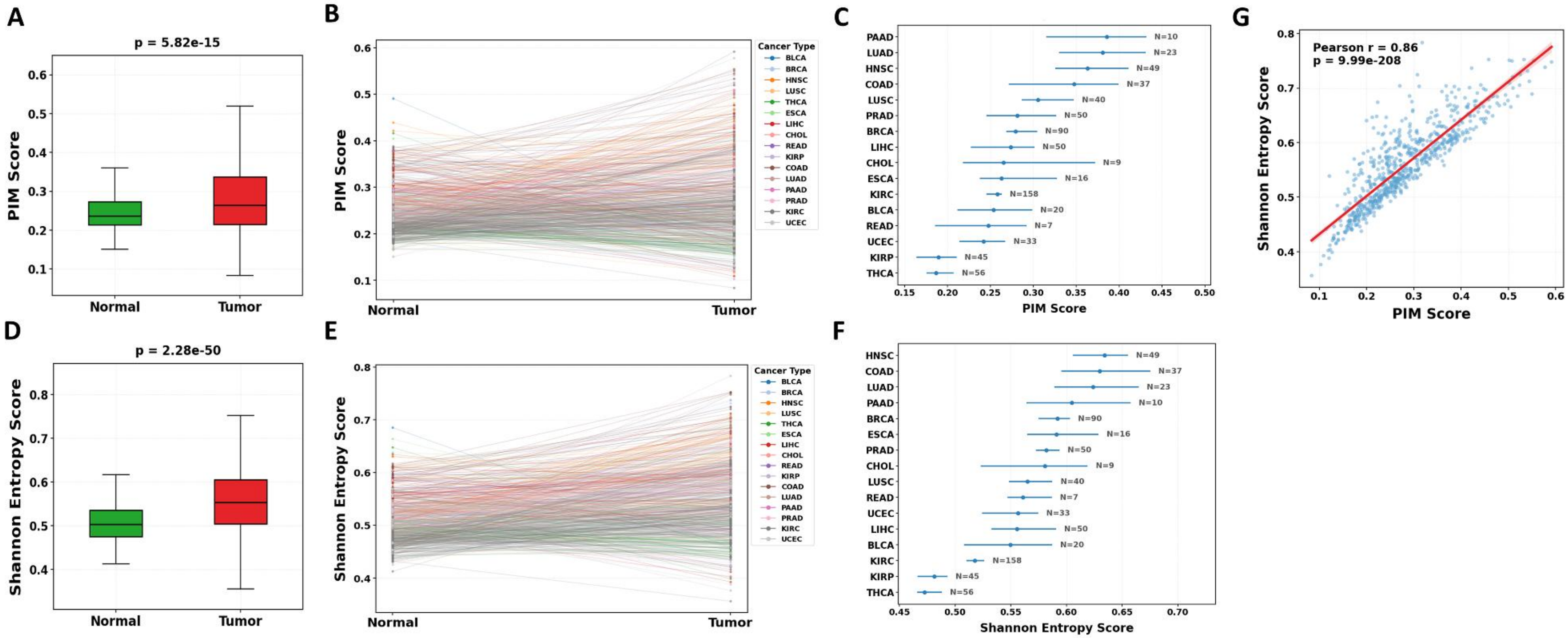


**Supplementary Figure 11. Pan-cancer comparison and concordance of PIM and Shannon entropy in matched normal and tumor tissues. A)** PIM scores in pooled matched normal and tumor samples. **B)** Paired PIM scores for individual normal–tumor pairs, with lines colored by cancer type. **C)** Cancer type–specific PIM score summaries; horizontal error bars indicate the corresponding uncertainty intervals and N denotes the number of matched pairs. **D)** Shannon entropy scores in pooled matched normal and tumor samples. **E)** Paired Shannon entropy scores for individual normal–tumor pairs, with lines colored by cancer type. **F)** Cancer type–specific Shannon entropy score summaries with corresponding error bars and numbers of matched pairs. **G)** Correlation between PIM and Shannon entropy scores across samples. The red line indicates the fitted linear regression; the Pearson correlation coefficient and corresponding *P* value are shown within the panel.

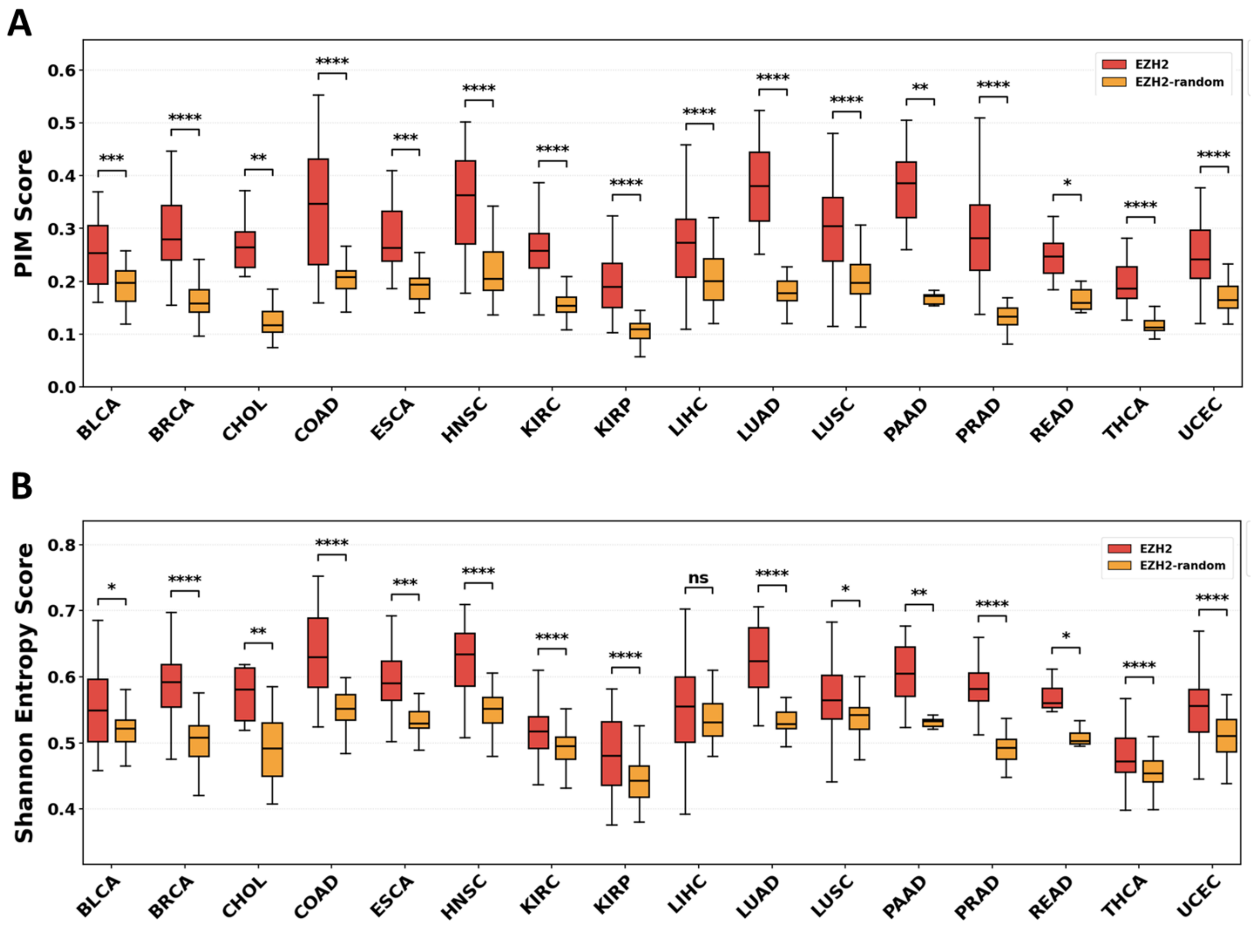


**Supplementary Figure 12. Comparison of DNA methylation variability between EZH2-binding regions and matched random genomic regions across cancer types. A)** PIM scores within EZH2-binding regions and corresponding matched random control regions across 16 cancer types. **B)** Shannon entropy scores within EZH2-binding regions and corresponding matched random control regions across the same cancer types. Boxplots show the median, interquartile range, and whiskers. Statistical significance is indicated above each EZH2 versus matched-random comparison (ns, not significant; *P* < 0.05; **P** < 0.01; ***P*** < 0.001; **P** < 0.0001).

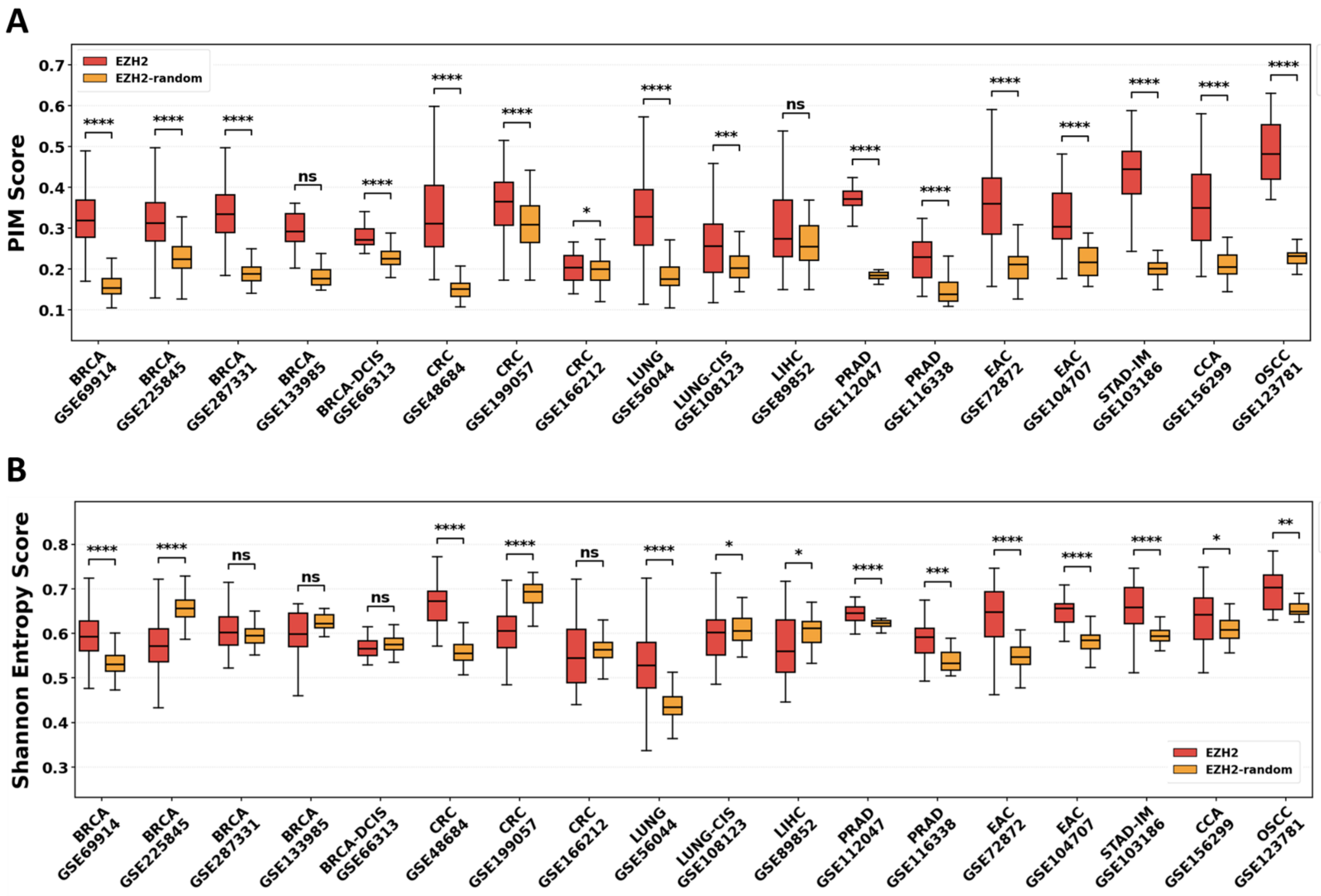


**Supplementary Figure 13. Comparison of DNA methylation variability between EZH2-binding regions and matched random regions across independent validation cohorts. A)** PIM scores within EZH2-binding regions and corresponding matched random control regions across the indicated GEO validation datasets and tissue types. **B)** Shannon entropy scores within EZH2-binding regions and corresponding matched random control regions across the same datasets. Boxplots show the median, interquartile range, and whiskers. Statistical significance is indicated above each comparison (ns, not significant; *P* < 0.05; **P** < 0.01; ***P*** < 0.001; **P** < 0.0001).

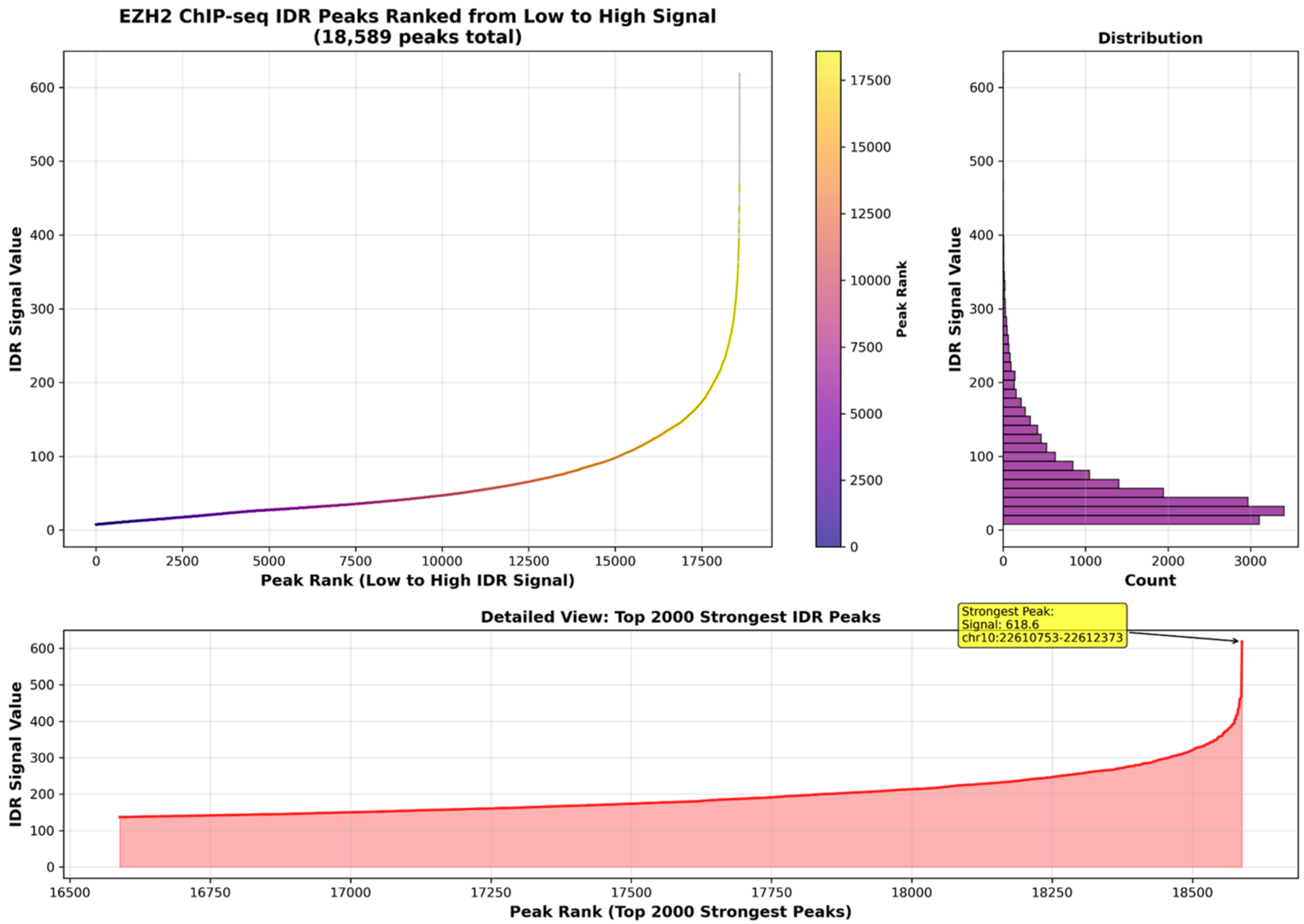


**Supplementary Figure 14. Distribution and ranking of EZH2 ChIP-seq IDR peak signal intensities.** EZH2 ChIP-seq irreproducible discovery rate (IDR) peaks were ranked from lowest to highest signal intensity across 18,589 peaks. **A)** Ranked distribution of EZH2 IDR signal values, with point color indicating peak rank. **B)** Histogram showing the distribution of EZH2 IDR signal values across all peaks. **C)** Detailed view of the 2,000 strongest EZH2 IDR peaks ranked by signal intensity. The highest-signal peak is annotated with its IDR signal value and genomic coordinates

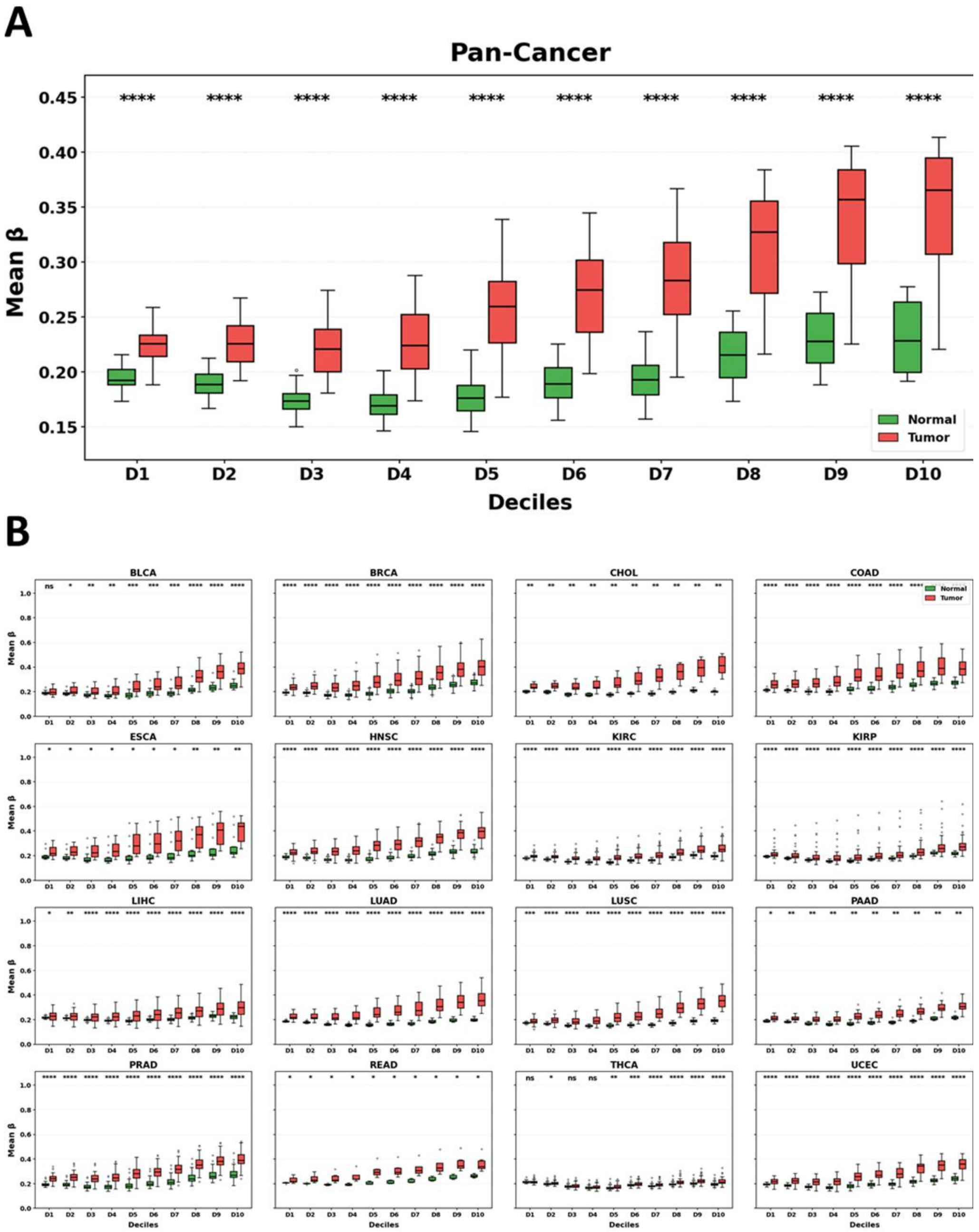


**Supplementary Figure 15. DNA methylation levels across EZH2-binding intensity deciles in matched normal and tumor tissues. A)** Pan-cancer comparison of mean DNA methylation (β value) across ten deciles (D1–D10) of EZH2-binding intensity in matched normal and tumor samples. **B)** Cancer type–specific comparisons of mean DNA methylation across EZH2-binding deciles for 16 cancer types. Normal tissues are shown in green and tumor tissues in red. Boxplots show the median, interquartile range, whiskers, and outliers. Statistical significance for tumor–normal comparisons within each decile is indicated above the corresponding groups (ns, not significant; *P* < 0.05; **P** < 0.01; ***P*** < 0.001; **P** < 0.0001).

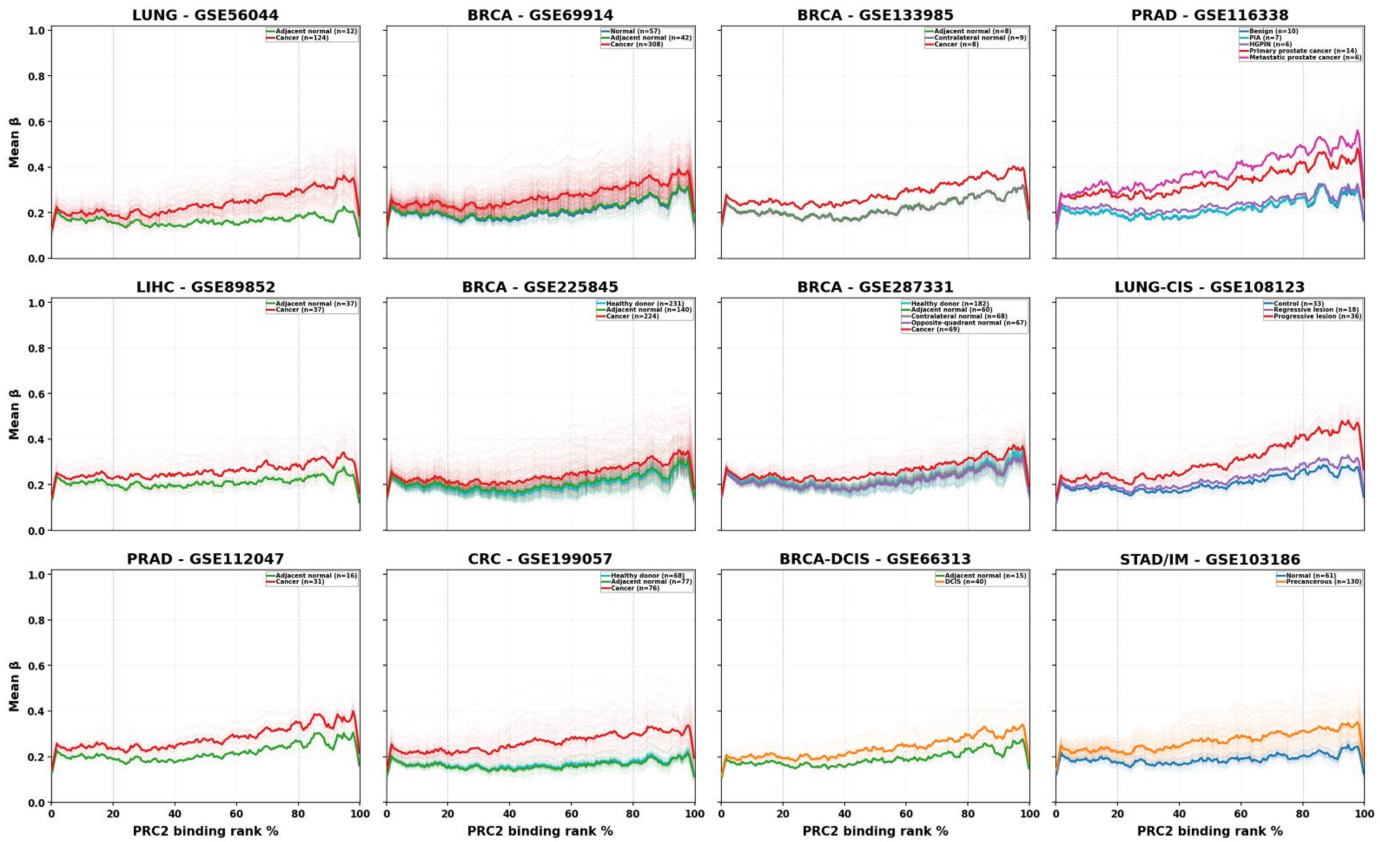


**Supplementary Figure 16. DNA methylation levels across increasing PRC2-binding intensity in independent validation cohorts.** Mean DNA methylation (β value) is plotted across genomic regions ranked from low to high PRC2 binding in 12 independent GEO datasets: lung cancer (GSE56044), breast cancer (GSE69914, GSE133985, GSE225845, and GSE287331), prostate cancer (GSE116338 and GSE112047), hepatocellular carcinoma (GSE89852), pre-invasive lung squamous lesions (GSE108123), colorectal cancer (GSE199057), breast ductal carcinoma in situ (GSE66313), and gastric intestinal metaplasia (GSE103186). Thin lines represent individual samples and solid lines represent group means. The x-axis indicates PRC2-binding rank percentile, and dashed vertical lines mark the 20th and 80th percentiles. Tissue and disease-state groups, together with corresponding sample numbers, are indicated within each panel.

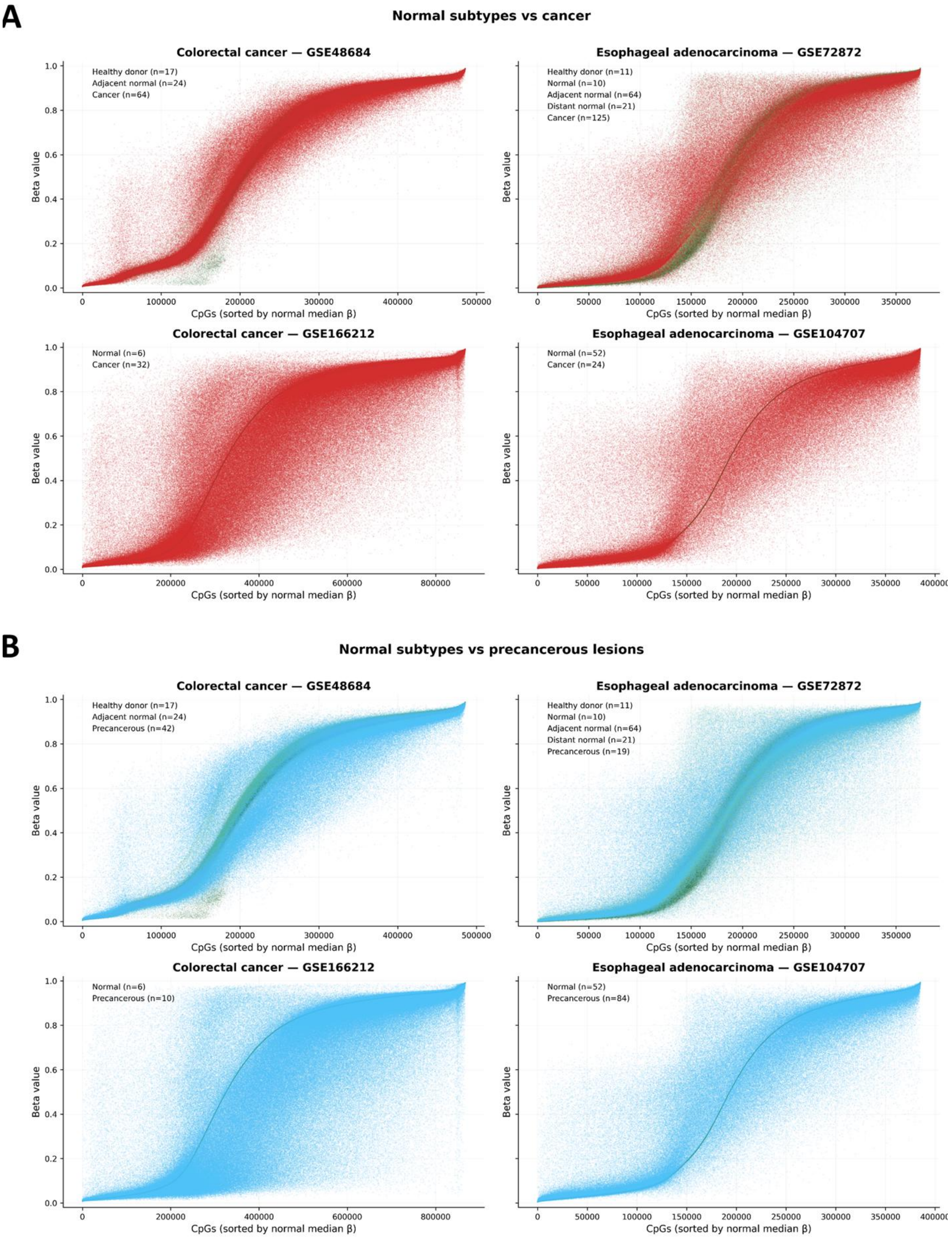
A
Normal subtypes vs cancer
Colorectal cancer — GSE48684
Healthy donor (n=17)
Adjacent normal (n=24)
Cancer (n=64)
Esophageal adenocarcinoma — GSE72872
Healthy donor (n=11)
Normal (n=10)
Adjacent normal (n=64)
Distant normal (n=21)
Cancer (n=125)
Colorectal cancer — GSE166212
Normal (n=6)
Cancer (n=32)
Esophageal adenocarcinoma — GSE104707
Normal (n=52)
Cancer (n=24)
Beta value
CpGs (sorted by normal median β)
B
Normal subtypes vs precancerous lesions
Colorectal cancer — GSE48684
Healthy donor (n=17)
Adjacent normal (n=24)
Precancerous (n=42)
Esophageal adenocarcinoma — GSE72872
Healthy donor (n=11)
Normal (n=10)
Adjacent normal (n=64)
Distant normal (n=21)
Precancerous (n=19)
Colorectal cancer — GSE166212
Normal (n=6)
Precancerous (n=10)
Esophageal adenocarcinoma — GSE104707
Normal (n=52)
Precancerous (n=84)
Beta value
CpGs (sorted by normal median β)

**Supplementary Figure 17. Genome-wide DNA methylation distributions across normal, precancerous, and cancerous tissues.** Genome-wide CpG methylation β values are plotted for independent colorectal cancer and esophageal adenocarcinoma cohorts, with CpGs ordered from low to high according to the median β value in the corresponding normal reference group. **A)** Comparison of healthy donor, normal, adjacent-normal, and distant-normal tissues with cancer samples in GSE48684, GSE72872, GSE166212, and GSE104707. **B)** Comparison of the corresponding normal tissue groups with precancerous samples in the same datasets. Individual sample β values are shown across the ordered CpGs; tissue groups and sample numbers are indicated within each panel.

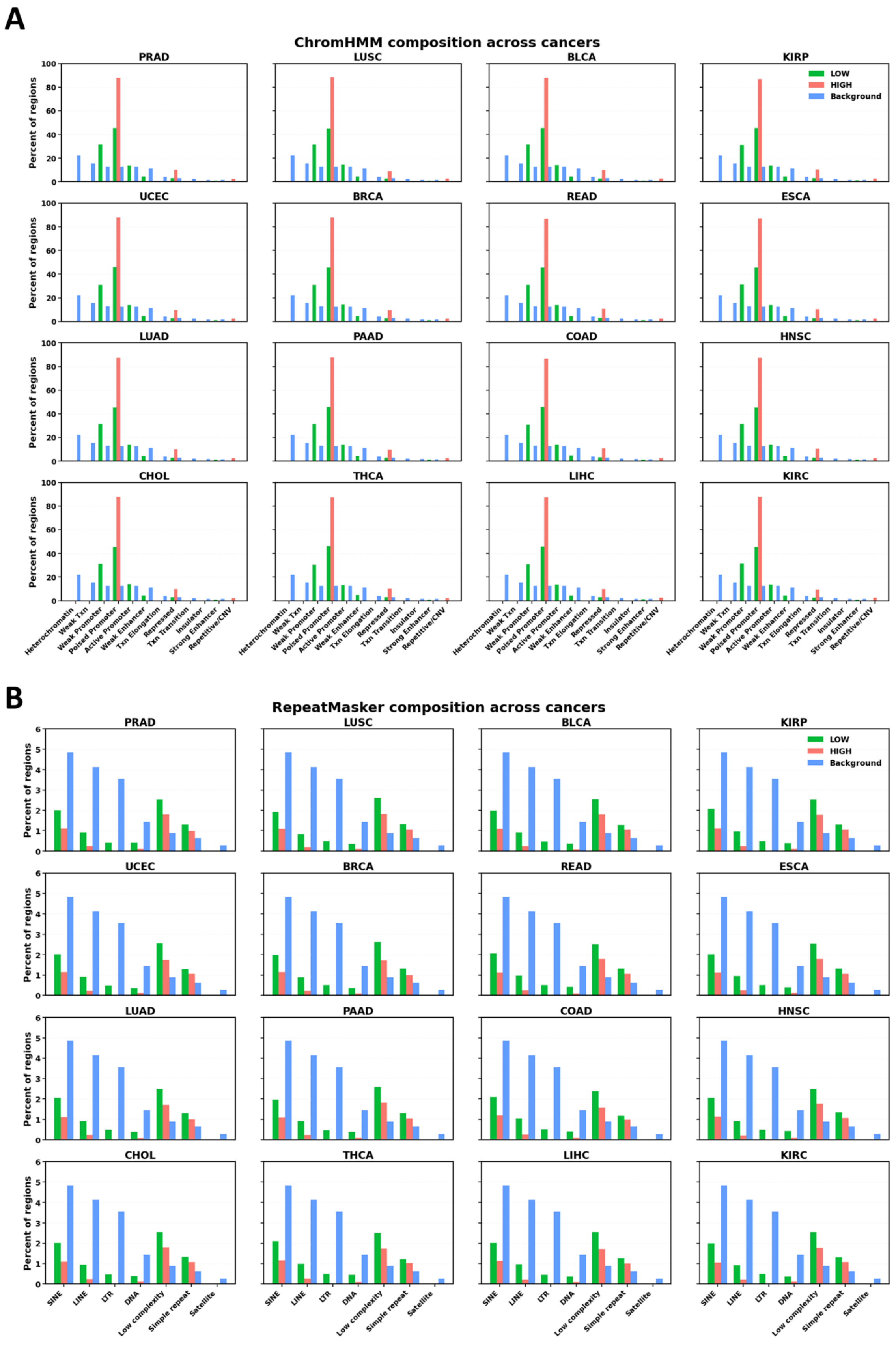
A
ChromHMM composition across cancers
PRAD
LUSC
BLCA
KIRP
UCEC
BRCA
READ
ESCA
LUAD
PAAD
COAD
HNSC
CHOL
THCA
LIHC
KIRC
LOW
HIGH
Background
Percent of regions
Heterochromatin
Weak Txn
Weak Promoter
Poised Promoter
Active Promoter
Weak Enhancer
Txn Elongation
Repressed
Txn Transition
Insulator
Strong Enhancer
Repetitive/CNV
B
RepeatMasker composition across cancers
SINE
LINE
LTR
DNA
Low complexity
Simple repeat
Satellite

**Supplementary Figure 18. Chromatin-state and repeat-element composition of low- and high-EZH2-binding regions across cancer types. A)** ChromHMM state composition of low-EZH2-binding regions (LOW), high-EZH2-binding regions (HIGH), and background regions across the indicated cancer types. Bars represent the percentage of regions assigned to each ChromHMM state. **B)** RepeatMasker annotation composition of LOW, HIGH, and background regions across the same cancer types. Bars represent the percentage of regions overlapping the indicated repeat classes, including SINEs, LINEs, LTRs, DNA transposons, low-complexity regions, simple repeats, and satellite repeats. LOW, HIGH, and background regions are shown in green, red, and blue, respectively.

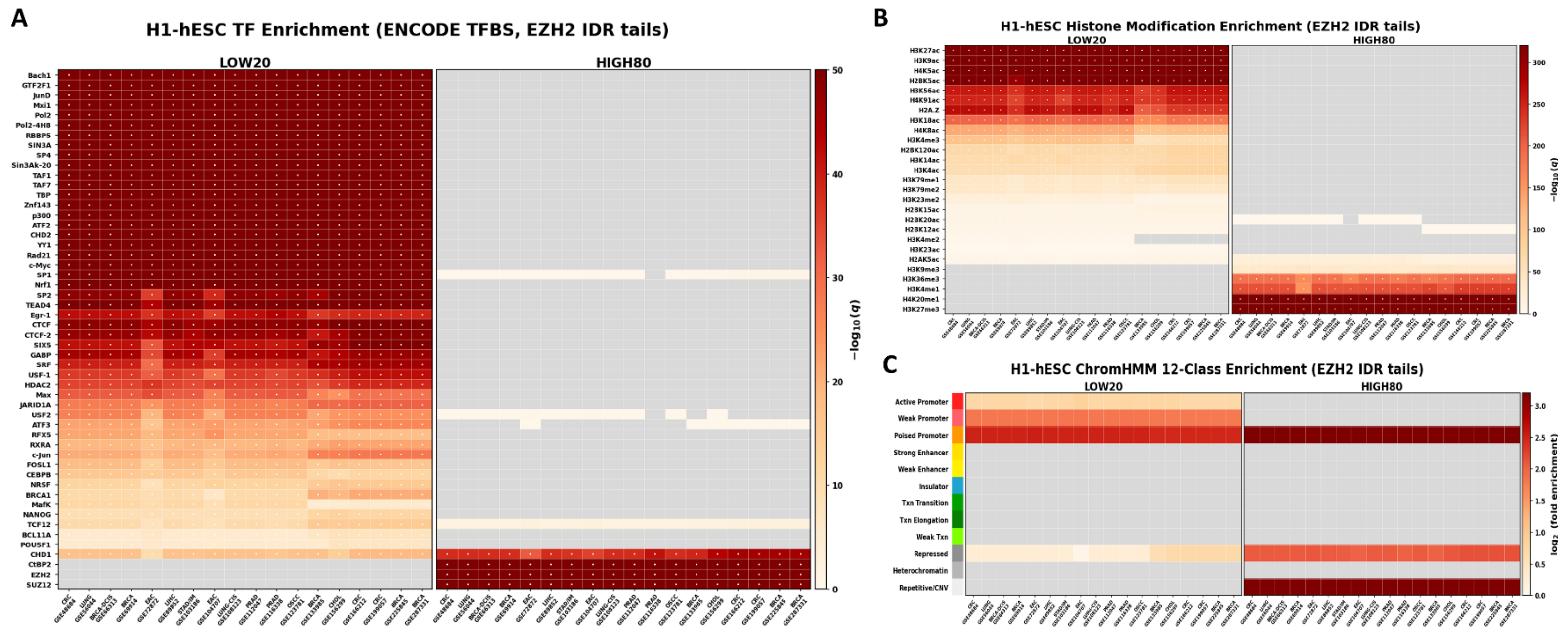


**Supplementary Figure 19. Regulatory and chromatin-state enrichment of low- and high-EZH2-binding regions across independent validation cohorts.** Genomic regions were stratified into the lowest 20% (LOW20) and highest 20% (HIGH80) of EZH2 ChIP-seq IDR signal and analyzed across the indicated GEO validation datasets using H1 human embryonic stem cell (H1-hESC) annotations. **A)** Enrichment of ENCODE transcription factor binding sites within LOW20 and HIGH80 EZH2-binding regions. **B)** Enrichment of histone modification annotations within LOW20 and HIGH80 regions. For **A)** and **B)**, color intensity represents −log10(*q*). **C)** Enrichment of H1-hESC ChromHMM chromatin-state annotations within LOW20 and HIGH80 EZH2-binding regions. Color intensity represents log2 fold enrichment. Gray cells indicate annotations without enrichment values in the corresponding comparison.

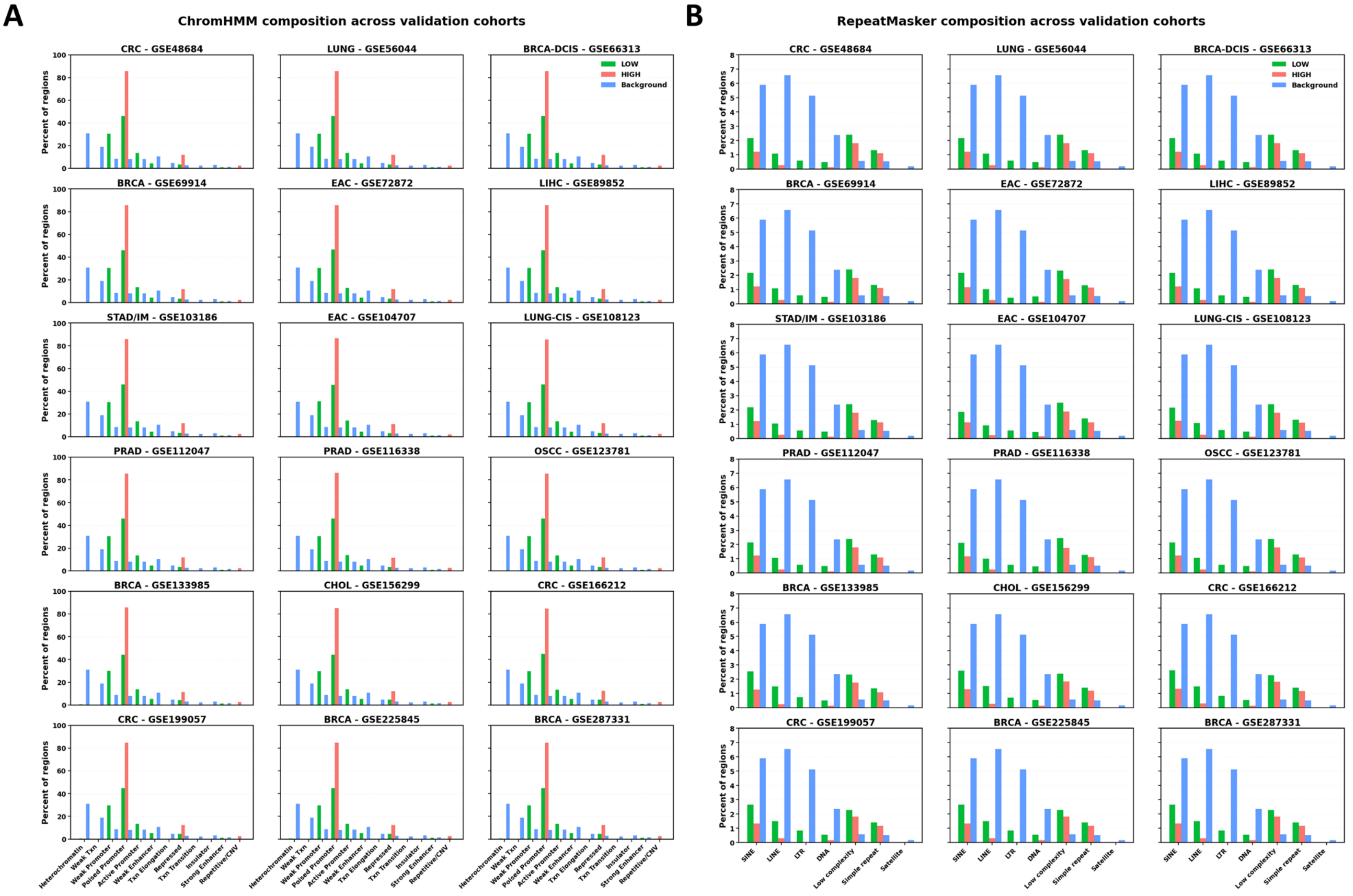
A
ChromHMM composition across validation cohorts
B
RepeatMasker composition across validation cohorts
CRC - GSE48684
LUNG - GSE56044
BRCA-DCIS - GSE66313
BRCA - GSE69914
EAC - GSE72872
LIHC - GSE89852
STAD/IM - GSE103186
EAC - GSE104707
LUNG-CIS - GSE108123
PRAD - GSE112047
PRAD - GSE116338
OSCC - GSE123781
BRCA - GSE133985
CHOL - GSE156299
CRC - GSE166212
CRC - GSE199057
BRCA - GSE225845
BRCA - GSE287331
LOW
HIGH
Background
Percent of regions
Heterochromatin
Weak Txn
Weak Promoter
Poised Promoter
Active Promoter
Weak Enhancer
Txn Elongation
Repressed
Txn Transition
Insulator
Strong Enhancer
Repetitive/CNV
SINE
LINE
LTR
DNA
Low complexity
Simple repeat
Satellite

**Supplementary Figure 20. Chromatin-state and repeat-element composition of low- and high-EZH2-binding regions across independent validation cohorts. A)** ChromHMM state composition of low-EZH2-binding regions (LOW), high-EZH2-binding regions (HIGH), and background regions across the indicated validation datasets. Bars represent the percentage of regions assigned to each ChromHMM state. **B)** RepeatMasker annotation composition of LOW, HIGH, and background regions across the same validation cohorts. Bars represent the percentage of regions overlapping the indicated repeat classes, including SINEs, LINEs, LTRs, DNA transposons, low-complexity regions, simple repeats, and satellite repeats. LOW, HIGH, and background regions are shown in green, red, and blue, respectively.

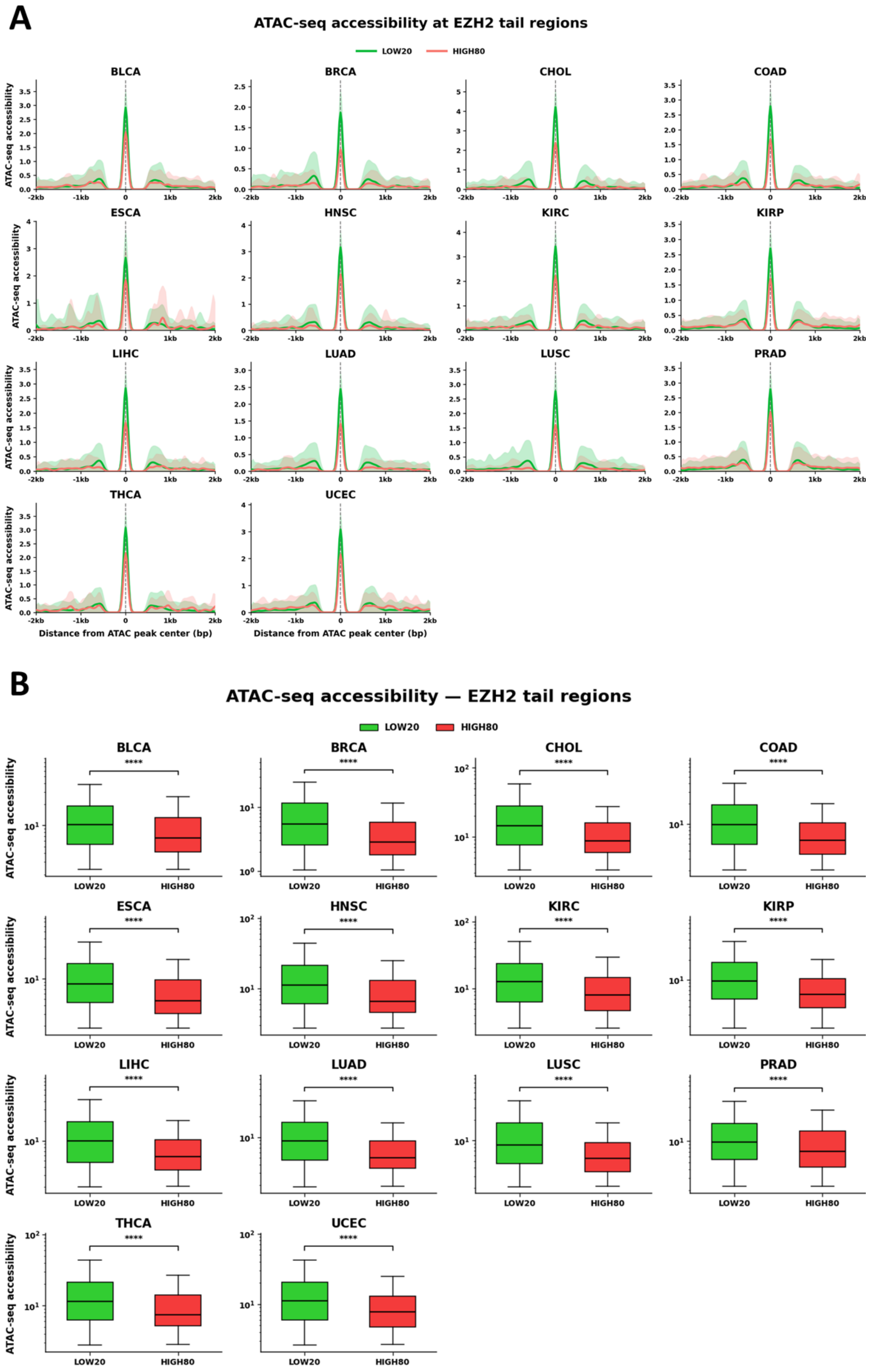
A
ATAC-seq accessibility at EZH2 tail regions
LOW20
HIGH80
BLCA
BRCA
CHOL
COAD
ESCA
HNSC
KIRC
KIRP
LIHC
LUAD
LUSC
PRAD
THCA
UCEC
ATAC-seq accessibility
Distance from ATAC peak center (bp)
B
ATAC-seq accessibility — EZH2 tail regions
LOW20
HIGH80
****

**Supplementary Figure 21. Chromatin accessibility across low- and high-EZH2-binding regions. A)** Aggregate ATAC-seq accessibility profiles centered on ATAC-seq peak summits within low-EZH2-binding regions (LOW20) and high-EZH2-binding regions (HIGH80) across the indicated cancer types. Profiles are shown across ±2 kb from the ATAC-seq peak center, with shaded areas indicating variability around the mean signal. **B)** Boxplot comparison of ATAC-seq accessibility between LOW20 and HIGH80 EZH2-binding regions for each cancer type. Boxplots show the median, interquartile range, and whiskers. Statistical significance is indicated above each LOW20 versus HIGH80 comparison (****P < 0.0001).

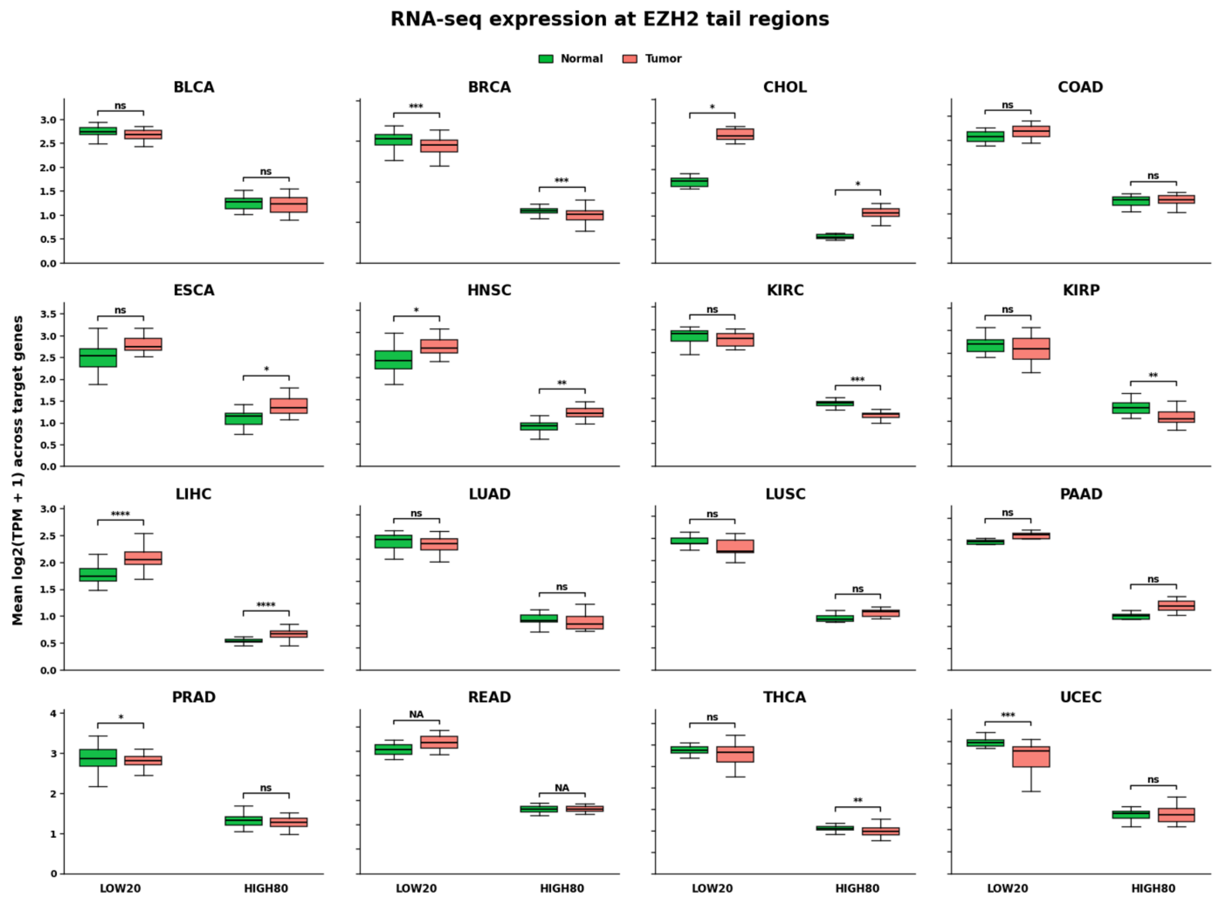


**Supplementary Figure 22. Gene expression across low- and high-EZH2-binding regions in normal and tumor tissues.** RNA-seq expression is shown for genes associated with the lowest 20% (LOW20) and highest 20% (HIGH80) EZH2-binding regions across the indicated cancer types. Expression is represented as the mean log2(TPM + 1) across target genes. Normal tissues are shown in green and tumor tissues in red. Boxplots show the median, interquartile range, and whiskers. Statistical significance for normal versus tumor comparisons within LOW20 and HIGH80 groups is indicated above the corresponding boxplots (ns, not significant; *P* < 0.05; **P** < 0.01; ***P*** < 0.001; **P** < 0.0001; NA, not available).

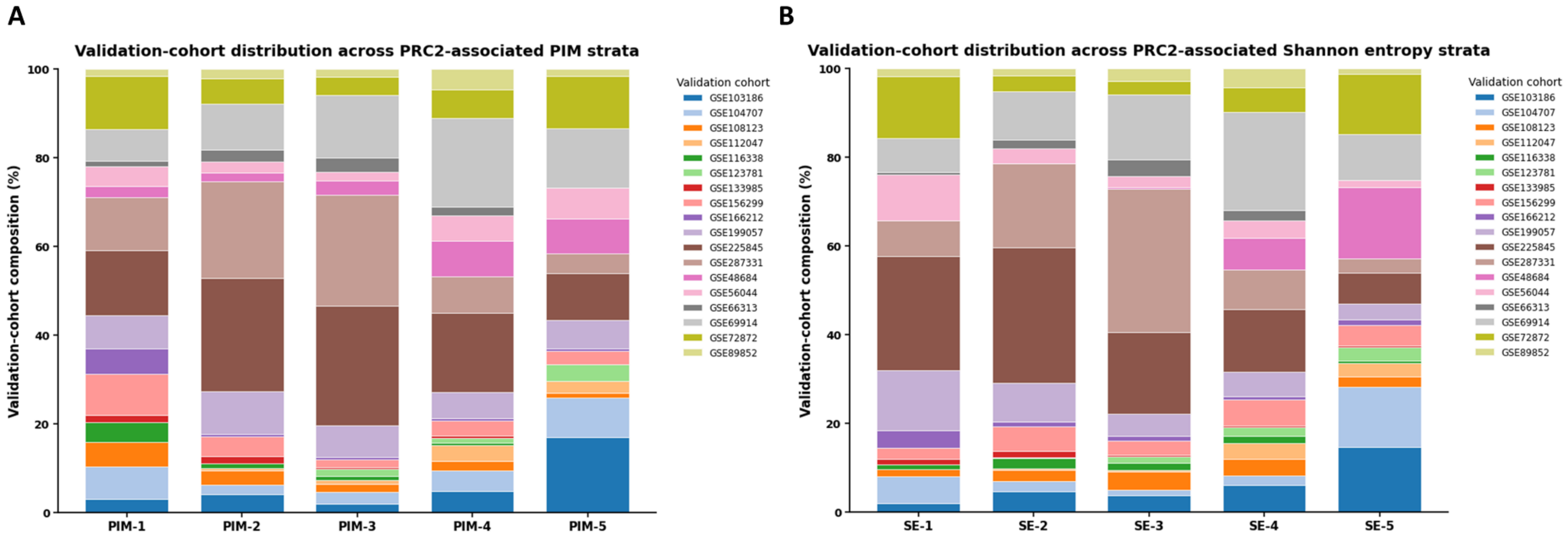


**Supplementary Figure 23. Distribution of validation cohorts across PRC2-associated DNA methylation disorder strata. A)** Relative contribution of individual validation cohorts across five strata of PRC2-associated PIM scores (PIM-1 to PIM-5). **B)** Relative contribution of individual validation cohorts across five strata of PRC2-associated Shannon entropy scores (SE-1 to SE-5). Stacked bars show the percentage composition of each stratum, with colors indicating the corresponding GEO validation cohorts.
